# Functional Determination of Calcium Binding Sites Required for the Activation of Inositol 1,4,5-trisphosphate receptors

**DOI:** 10.1101/2022.03.07.482538

**Authors:** Vikas Arige, Lara E. Terry, Larry E. Wagner, Mariah R. Baker, Guizhen Fan, Irina I. Serysheva, David I. Yule

## Abstract

Inositol 1,4,5-trisphosphate (IP_3_) receptors (IP_3_Rs) initiate a diverse array of physiological responses by carefully orchestrating intracellular calcium (Ca^2+^) signals in response to various external cues. Notably, IP_3_R channel activity is determined by several obligatory factors including IP_3_, Ca^2+^ and ATP. The critical basic amino acid residues in the N-terminal IP_3_-binding core (IBC) region that facilitate IP_3_ binding are well characterized. In contrast, the residues conferring the biphasic regulation by Ca^2+^ are yet to be ascertained. Using comparative structural analysis of Ca^2+^ binding sites identified in two main families of intracellular Ca^2+^-release channels, ryanodine receptors (RyRs) and IP_3_Rs, we identified putative acidic residues coordinating Ca^2+^ in the cytosolic calcium sensor region in IP_3_Rs. We determined the consequences of substituting putative Ca^2+^ binding, acidic residues in IP_3_R family members. We show that the agonist-induced Ca^2+^ release, single channel open probability (P_0_) and Ca^2+^ sensitivities are markedly altered when the negative charge on the conserved acidic side chain residues are neutralized. Remarkably, neutralizing the negatively charged side chain on two of the residues individually in the putative Ca^2+^ binding pocket shifted the Ca^2+^ required to activate IP_3_R to higher concentrations, indicating that these residues likely are a component of the Ca^2+^ activation site in IP_3_R. Taken together, our findings indicate that Ca^2+^ binding to a well conserved activation site is a common underlying mechanism resulted in increased channel activity shared by IP_3_Rs and RyRs.

## Introduction

A rise in the intracellular calcium concentration [Ca^2+^]_i_ is essential for innumerable fundamental biological processes. Following stimulation, the concerted action of a diverse array of proteins, collectively termed the Ca^2+^ signaling “tool-kit”, functions to increase and sequester Ca^2+^ to precisely define the spatiotemporal characteristics of Ca^2+^ signals necessary to control diverse physiological endpoints with specificity and fidelity (1, 2). As a function of exquisite regulation of their activity by a multitude of diverse modulatory inputs, the inositol 1,4,5-trisphosphate receptor family (IP_3_R) of Ca^2+^ release channels are a central component of this cellular machinery. In vertebrates, there are three IP_3_R sub-types: IP_3_R Type 1 (IP_3_R1), IP_3_R Type 2 (IP_3_R2) and IP_3_R Type 3 (IP_3_R3) that share about 60-70% amino acid sequence identity (3-7). IP_3_Rs are ubiquitously expressed and predominantly localized to the endoplasmic reticulum (ER) membrane where functionally they are assembled as either homo- or hetero-tetramers. While binding of IP_3_ – generated as a result of activation of phospholipase C (PLC) – is necessary to gate the channel, it is also clear that the binding of Ca^2+^ itself is obligatory to activate the receptor (8). Thus, IP_3_ and Ca^2+^ are considered co-agonists of IP_3_R (9).

Early work using truncated IP_3_R1 established that the N-terminal 788 amino acid residues were required to bind IP_3_. Subsequently, mutagenesis approaches established the minimal IP_3_-binding core (IBC, residues 224-557) and identified the positively charged residues required for coordinating the phosphate moieties in IP_3_ (10). More recent X-ray crystallography of the isolated IBC domains (11, 12) and single-particle cryogenic electron microscopy (cryo-EM) studies (13-16) have revealed the conserved 3D architecture of the ligand binding domains (LBDs) that includes β-trefoil domains, β-TF1 (residues 1-225) and βTF2 (residues 226-435) and a helical armadillo repeat, ARM1 domain (residues 436-665), providing structural determinants for coordination of IP_3_ and adenophostin A, a structural mimetic of IP_3_, in the IP_3_-binding pocket. These studies showed that the ligand binding is accompanied by the conformational changes not only in the LBDs (11, 12, 14, 15), but involves global conformational changes in the cytoplasmic scaffold connected to the channel pore (14, 15). Critically, the functional relevance of these residues has also been confirmed by mutagenesis (17). Indeed, one of the residues (R269) important for IP_3_ binding is the locus of a point mutation (R269W) that results in spino-cerebellar ataxia in patients as a result of a poorly functional IP_3_R1 (18). Furthermore, mutagenesis of these residues, in increasing numbers of IP_3_R monomers within a concatenated tetramer has led to the demonstration that IP_3_ binding to all four subunits in an IP_3_R tetramer is an absolute pre-requisite for channel opening (19).

Extensive functional characterization at the level of Ca^2+^ release and at the single channel level have demonstrated that Ca^2+^ modulates IP_3_R channel opening in a biphasic manner (8, 20-25). In the presence of IP_3_, low concentrations of Ca^2+^ (<300 nM) facilitate channel opening, however, at higher concentrations Ca^2+^ promotes channel closure to result in a bell-shaped regulation of IP_3_R activity (8). These data are consistent with Ca^2+^ interacting at two distinct sites, one which results in activation and a further site which attenuates activity (26). Models of this regulation have postulated that IP_3_ binding modulates the affinity of the Ca^2+^ binding sites for Ca^2+^ to favor increased or decreased IP_3_R channel activity (24, 27-30). This control of IP_3_R activity by the interplay between co-agonists is widely considered to mechanistically underlie the hierarchy of Ca^2+^ signaling events observed in cells including localized Ca^2+^ blips, Ca^2+^ puffs, propagating Ca^2+^ waves and Ca^2+^ oscillations (21, 31-33).

While the determinants of IP_3_ binding are firmly established (10), elucidating the motifs that coordinate Ca^2+^ in IP_3_R have proved elusive and are yet to be functionally determined (34). Notably using gel overlay assays, several linear fragments of IP_3_R across various domains were shown to bind ^45^Ca^2+^ (35, 36). Nevertheless, point mutations at conserved acidic amino acids in regions identified in the IBC were without any effect on Ca^2+^ dependent activation of IP_3_R, ruling out the possibility of Ca^2+^ binding site/s in this domain (37). A further series of studies proposed glutamate (E) 2100 as a key residue in IP_3_R for activation by Ca^2+^. Mutating this residue diminished Ca^2+^ sensitivity for activation of IP_3_R1 by several fold, but also altered the IP_3_R sensitivity to inhibition by Ca^2+^ (38, 39). Nevertheless, there is little structural evidence supporting direct binding of this residue to Ca^2+^, indicating that this site perhaps has an allosteric role in stabilizing a confirmation favoring Ca^2+^ binding to bonafide Ca^2+^ binding site/s (21, 27).

Perhaps the best clues to potential Ca^2+^ binding sites in IP_3_R come from structural studies of the ryanodine receptor (RyR) (38-40). IP_3_Rs share structural and functional features with RyRs; both are intracellular Ca^2+^ release channels with tetrameric architecture and are regulated in a biphasic manner by Ca^2+^ (8). The cryo-EM structure of RyR determined in the presence and absence of its activating ligands (Ca^2+^/ATP/caffeine) revealed several regions which coordinate Ca^2+^ ions in RyRs. These include paired EF-hand domains in the central region of RyR and a pocket of negatively charged residues in the core solenoid and C-Terminal Domain (40). While mutation of the EF-hands did not alter Ca^2+^ activation of RYR2, mutation of the putative binding pocket in RyR1/2 altered Ca^2+^-dependent activation of RyRs (41, 42). Of note, an analogous Ca^2+^ binding site is present in the juxta membrane domain of IP_3_R between the third armadillo repeat (ARM3) and linker (LNK) domains and is conserved in IP_3_Rs across species and through IP_3_R subtypes (43). Recent cryo-EM analysis of human IP_3_R3 in the presence of IP_3_ and supra-physiological Ca^2+^ suggested two Ca^2+^ binding sites including a site within the putative Ca^2+^ sensor regions located at the interface between the ARM3 (residues 1587-2119) and LNK (residues 2538-2608) domains (15). However, the limited resolution in this structure precludes identification of any substantive interactions within the binding pockets. Moreover, the channel pore was not engaged under the aforementioned conditions, therefore, it is unclear whether either site relates to stimulation or inhibition of the channel.

In this study, based on our single-particle cryo-EM structure of IP_3_R1 determined at ∼3.2 Å resolution, a series of experiments were performed to examine the consequences of substituting putative Ca^2+^ binding conserved residues in IP_3_R sub-types. We generated stable cell lines expressing wild-type (WT) IP_3_Rs and compared agonist-evoked Ca^2+^ release, Ca^2+^ puffs and single channel properties with their counterparts harboring substitutions at putative Ca^2+^ coordinating sites. Neutralizing the negative charge on side chain residues at these conserved Ca^2+^ coordinating residues decimated both Ca^2+^ release and Ca^2+^ puff activity evoked by IP_3_ generating agonists. Remarkably, substitution with a smaller, but similarly charged, residue markedly diminished agonist-induced Ca^2+^ release and Ca^2+^ puffs compared to their WT counterparts. Of note, comparison of single channel properties revealed that the Ca^2+^ sensitivity was dramatically right shifted by charge neutralization, indicating that this site is likely the Ca^2+^ activation site. In contrast, retaining charge on the side chain residue had little effect on Ca^2+^ sensitivity, however, open probability of channel was significantly diminished. Overall, our investigations demonstrate a critical role for the negative charge on side chains of conserved Ca^2+^ coordinating residues in IP_3_Rs for electrostatic interactions with Ca^2+^ to facilitate channel opening and subsequent, agonist-induced Ca^2+^ release.

## Results

### Identification of putative Ca^2+^ coordinating residues in IP_3_Rs

Based on our comparative structural analysis of RyR and IP_3_R channels, the three amino acid residues of human IP_3_R1 (hIP_3_R1) (NP_001093422.2), E2002, E1938 and T2614 from the putative Ca^2+^ binding pocket of hIP_3_R1 are highly conserved across various organisms in both IP_3_R (15) and RyR sub-types (40, 41, 43) (Fig. 1A-E and Fig. S1A). In contrast, the putative Ca^2+^ binding pocket located between α-helical domain (HD) and the second armadillo repeat (ARM2) in hIP_3_R3 (15) are not conserved between RyRs and IP_3_Rs (Fig. S1B). These observations indicate that conserved residues forming the Ca^2+^ binding site in two main families of intracellular Ca^2+^-release channels are good candidates to mediate regulation of IP_3_R by Ca^2+^. This prediction has been confirmed by our concurrent cryo-EM study of IP_3_R1 in Ca^2+^ bound states (Fig. 1A-D).

**Figure 1.**
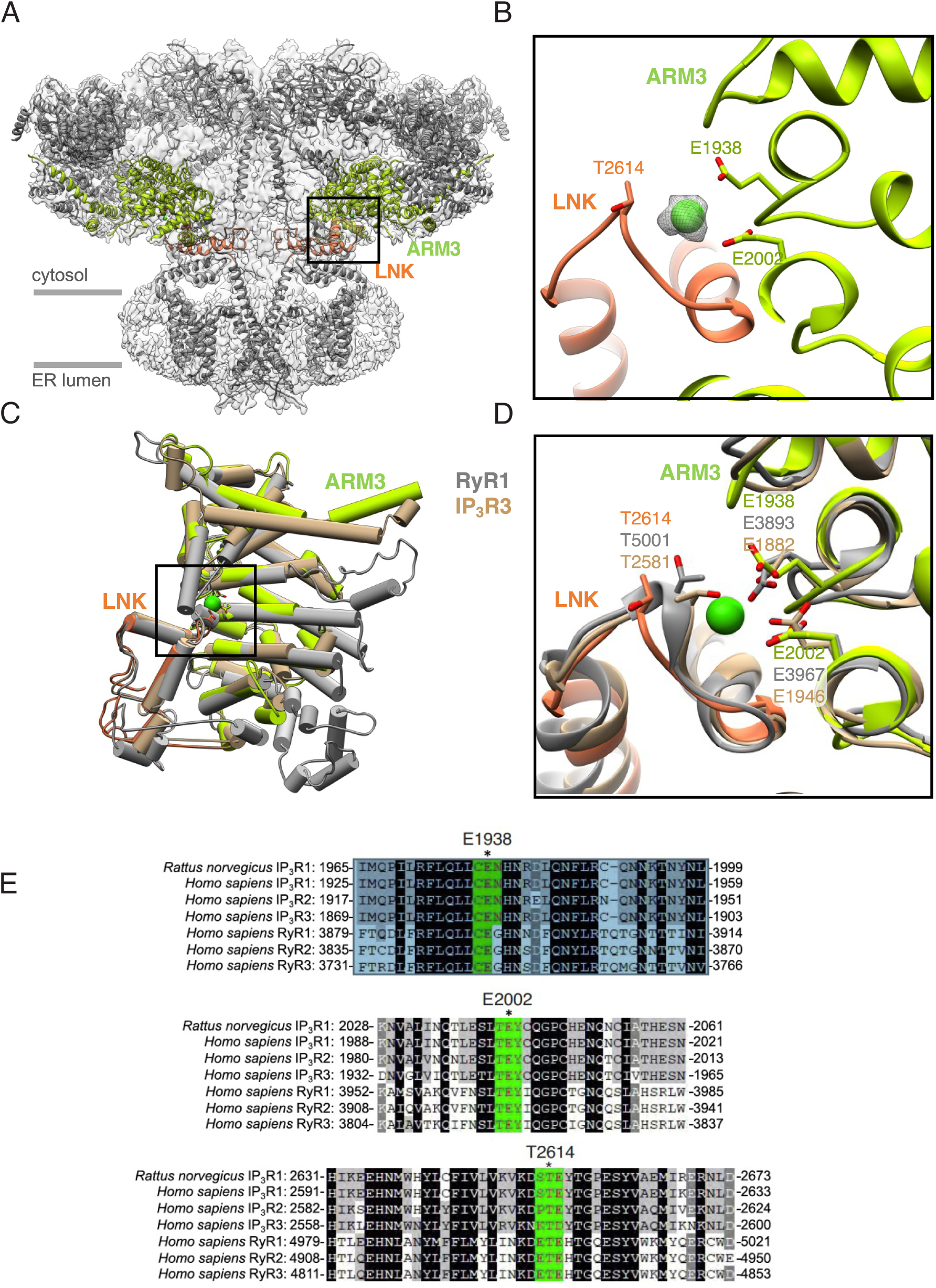
Structural and sequence conservation of the Ca^2+^ binding site across IP_3_R and RyR families of Ca^2+^ release channels. **A**. 3D structure of IP_3_R1 determined in the presence of 50 μM Ca^2+^ at ∼3.2 Å resolution by single-particle cryo-EM: a 3D model of the channel is overlayed with the corresponding cryo-EM density map of the tetrameric IP_3_R1-lipid nanodisc complex; shown are two opposing subunits with LNK and ARM3 domains color-coded; viewed along the membrane plane. **B**. Close-up view of Ca^2+^ binding site formed by LNK (orange) and ARM3 (chartreuse) indicated by the gray dashed line in ‘A’ panel; conserved residues in the Ca^2+^ binding pocket are labeled; the density corresponding to Ca^2+^ is depicted as gray mesh, contoured at 5σ. **C**. 3D structural conservation of the Ca^2+^ sensor region in IP_3_R1 (as color coded in ‘A’), IP_3_R3 (PDB ID: 2DR2) and RyR1 (PDB ID: 5T15). Cα RMS deviation is 1.67 Å. **D**. Overlay of IP_3_R1, IP_3_R3 and RyR1 Ca^2+^ binding pockets with identical residues labeled. **E**. Sequence alignment for the Ca^2+^-sensor region across all three IP_3_R and RyR sub-types. The conserved E1938, E2002 and T2614 residues in IP_3_R1 are highlighted in green and indicated by asterisks.

### Substitutions at the putative Ca^2+^ coordinating residues in hIP_3_R1 diminished agonist-induced Ca^2+^ release

IP_3_R1 is the best studied subtype in terms of regulation by Ca^2+^ (13, 44, 45). Therefore, we stably over-expressed WT hIP_3_R1 or hIP_3_R1 with substitutions at the 2002 glutamic acid (E) residue (Fig. 1B) to aspartic acid (D), alanine (A), or glutamine (Q) in HEK-3KO cells previously generated in our laboratory by CRISPR/Cas9 technology to lack native IP_3_Rs (19). The rationale for the substitutions chosen being that the E-D residue change retained side chain charge; the E-Q substitution results in a similar sized side chain that does not retain side chain charge but is still hydrophilic and the E-A substitution retains the β carbon moiety but has no other side chain chemistry.

When expressed in HEK-3KO cells, IP_3_R1 protein levels of exogenously (exo) expressed WT hIP_3_R1, hIP_3_R1 E2002D, hIP_3_R1 E2002A and hIP_3_R1 E2002Q stable cell lines were higher than WT HEK293 and endogenous (endo) hIP_3_R1 cells, previously generated in our laboratory by CRISPR/Cas9 technology to lack the other two native IP_3_Rs (19) (Fig. 2A, B). To investigate the functional consequences of these substitutions on hIP_3_R1 channel activity, we first performed single cell imaging assays to assess IP_3_-induced Ca^2+^ release using carbachol (CCh) and trypsin as agonists acting via the G_α_q-coupled M3 muscarinic receptor and Protease-Activated Receptor 2 (PAR2), respectively. CCh-induced Ca^2+^ release was markedly diminished in the hIP_3_R1 E2002D cells when compared to exo hIP_3_R1 cells at lower doses of CCh (3, 10 µM), however, such differences disappeared at a higher dose of CCh (30 µM) presumably due to recruitment of additional IP_3_Rs due to Ca^2+^-induced Ca^2+^ release (CICR) at higher concentrations of CCh (Fig. 2C, D). CCh-induced Ca^2+^ release was significantly attenuated in hIP_3_R1 E2002A and hIP_3_R1 E2002Q cells as compared to exo hIP_3_R1 cells at all doses of CCh (3, 10, 30 µM) and to endo hIP_3_R1 cells at a higher dose of CCh (30 µM) (Fig. 2C, D). Moreover, the percentage of responding cells were also significantly lower in stable cells with substitutions at the E2002 residue compared to WT cells (Fig. S2A). Similarly, trypsin-evoked Ca^2+^ release was also significantly attenuated in cells with substitutions at the E2002 site when compared to both endo and exo hIP_3_R1 cells (Fig. S2C, D). The HEK-3KO cells, as previously demonstrated, failed to respond to stimulation with either trypsin (Fig. S2C, D) (18, 19) or CCh (Fig. 2C-E). Further, in a population-based assay performed on a FlexStation3 plate reader with microfluidics, CCh-induced Ca^2+^ release as reflected by both the maximum amplitude changes (Fig. 2E) and area under the curve (AUC) (Fig. S2B) were markedly attenuated in E2002D and decimated in E2002A and E2002Q cells when compared to both exo and endo IP_3_R1 cells. These results suggest that the negative charge on the side chain residue at 2002 position in hIP_3_R1 is critical for binding to Ca^2+^ and neutralizing this charge prevented IP_3_-induced Ca^2+^ release and IP_3_R1 activation.

**Figure 2.**
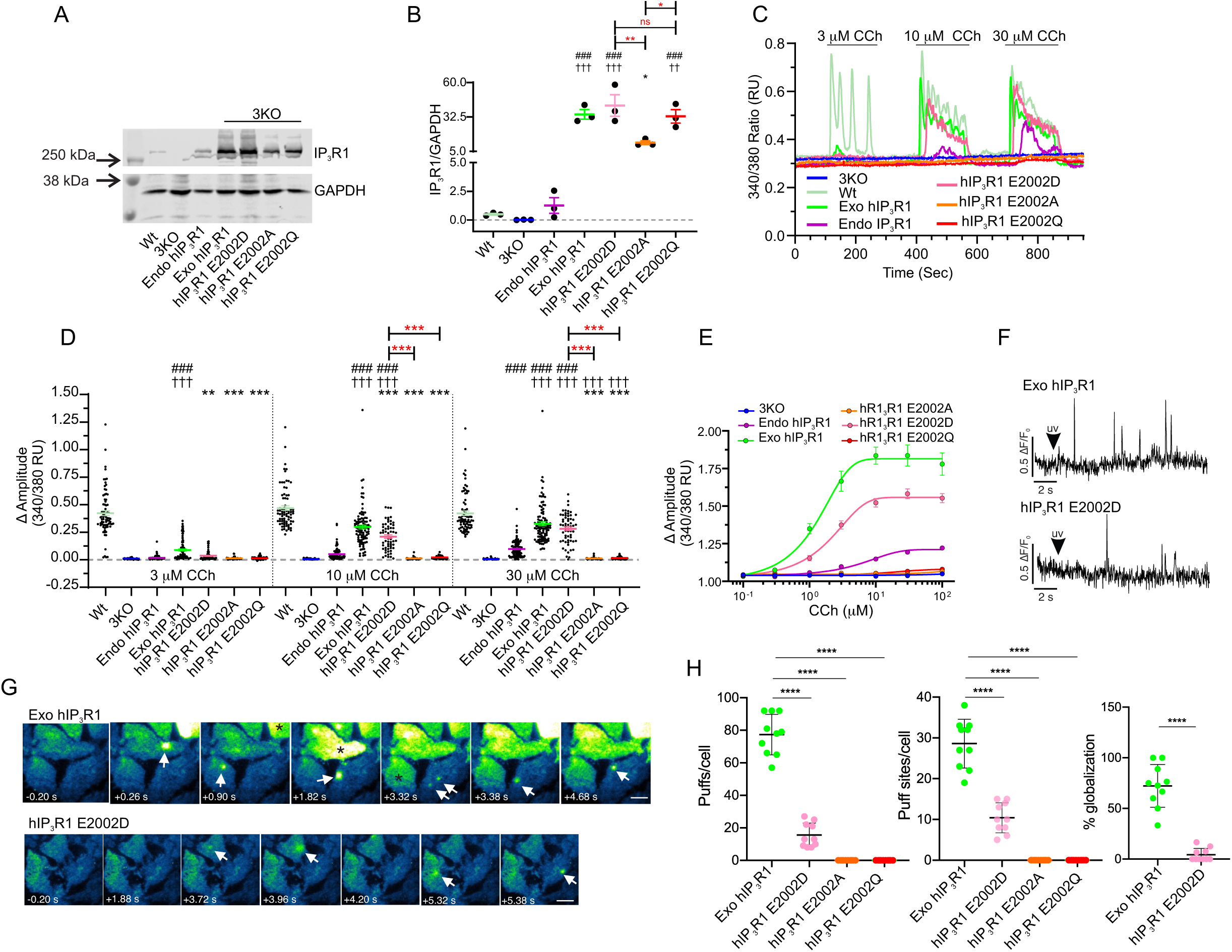
Substituting E2002 with aspartate (D), alanine (A) or glutamine (Q) residue in hIP_3_R1 significantly diminished agonist-induced Ca^2+^ signals when stably expressed in HEK-3KO cells. Stable hIP_3_R1 cell lines with a substitution at E2002 position to D, A or Q were generated in HEK-3KO cells. **A**. A representative western blot depicting IP_3_R1 and GAPDH protein levels in wild-type HEK293 (WT), HEK-3KO (3KO), HEK-2KO (lacking native IP_3_R2 and IP_3_R3 sub-types, endo hIP_3_R1), 3KO cells over-expressing WT IP_3_R1 (exo hIP_3_R1) and E2002D/A/Q substitution (hR1 E2002D/A/Q). **B**. Scatter plot depicting quantification of IP_3_R1 protein level normalized to GAPDH from three-independent western blots. **C**. Representative traces showing CCh-induced (3, 10, 30 µM) Ca^2+^-release from 3KO (blue), Wt (pale yellow-green), endo hIP_3_R1 (purple), exo hIP_3_R1 (green), hIP_3_R1 E2002D (pink), hIP_3_R1 E2002A (orange), hIP_3_R1 E2002Q (red) cells in single cell assays. **D**. Scatter plot summarizing change in amplitude (Peak ratio – Basal ratio: average of 20 ratio points immediately preceding addition) to increasing doses of CCh (3, 10, 30 µM) for experiments similar to those shown in C. **E**. Dose-response curve showing maximum amplitude from Fura-2/AM loaded 3KO (blue), endo hIP_3_R1 (purple), exo hIP_3_R1 (green), hIP_3_R1 E2002D (pink), hIP_3_R1 E2002A (orange) and hIP_3_R1 E2002Q (red) cells in population-based assays when treated with increasing concentrations (100 nM, 300 nM, 1 μM, 3 μM, 10 μM, 30 μM and 100 μM) of CCh. **F**. Representative traces of Cal-520 fluorescence ratios (ΔF/F_0_) from the center of a single puff site (∼1 × 1 µm) evoked by photolysis of ci-IP_3_ at 2 sec (indicated by arrow) in exo hIP_3_R1 (upper trace) and hIP_3_R1 E2002D (lower trace) cells using a TIRF microscope. **G**. TIRF images from exo hIP_3_R1 (upper panel) and hIP_3_R1 E2002D (lower panel) showing Ca^2+^ puffs at indicated time point. Typical of at least 10 independent experiments. Arrows indicate Ca^2+^ puffs/puff sites, asterisks indicate globalization of Ca^2+^ signals. **H**. Scatter plots summarizing number of puffs/cell, puff sites/cell and % globalization in exo hIP_3_R1 cells (green), hIP_3_R1 E2002D (pink), hIP_3_R1 E2002A (orange) and hIP_3_R1 E2002Q (red) cells (N=10 cells). Data are mean ± SEM of three (N = 3) independent experiments. ###*P* < 0.001 when compared to 3KO cell line, ††*P* < 0.01, †††*P* < 0.001 when compared to endo hIP_3_R1 cell line and \**P* < 0.05, \*\**P* < 0.01, \*\*\**P* < 0.001, \*\*\*\**P* < 0.0001 when compared to exo hIP_3_R1 cell line; one-way ANOVA with Tukey’s test was performed. Unless otherwise stated, all data above comes from at least N=3 experiments.

Next, to measure the activity of the hIP_3_R more directly and without the global effects on neighboring IP_3_R by increasing Ca^2+^, we investigated the effect of substitutions at the E2002 site on the fundamental Ca^2+^ signals-mediated by IP_3_Rs (termed Ca^2+^ puffs) upon uncaging ci-IP_3_ using TIRF microscopy (31, 33, 46-48). As shown in the representative traces (Fig. 2F) and images (Fig. 2G), the number of fundamental Ca^2+^ signals obtained from the hIP_3_R1 E2002D cells were significantly diminished when compared to WT exo hIP_3_R1 cells following uncaging of ci-IP_3._ The number of puffs per cell, puff sites per cell and the percentage of cells in which Ca^2+^ signals globalized were significantly attenuated in hIP_3_R1 E2002D cells as compared to WT hIP_3_R1 cells (Fig. 2H). The rise and fall times did not differ between these two cells (Fig. S2E). We failed to detect any puffs in cells expressing hIP_3_R1 with either E2002A and E2002Q, again reinforcing the importance of the negative charge at this residue (Fig. 2H).

### Substituting the putative Ca^2+^ coordinating residue in hIP_3_R3 diminished agonist-induced Ca^2+^ release

We postulated that if E2002 indeed represented an important charged residue for the co-ordination of Ca^2+^ and subsequent activity of hIP_3_R1, then the functional effect of mutating this residue should be conserved in other hIP_3_R subtypes. Therefore, we generated stable cell lines in HEK-3KO with substitution at the analogous E1946 residue in hIP_3_R3 (Fig. 1E). The IP_3_R3 protein levels of the hIP_3_R3 E1946Q stable cell line were slightly more than the WT HEK293 and endogenous hIP_3_R3 cells, previously generated in our laboratory by CRISPR/Cas9 technology to lack the other two native IP_3_Rs (Fig. 3A, B). To determine the functional consequence of this substitution, as before, we performed single cell imaging assays to assess IP_3_-induced Ca^2+^ release using CCh and trypsin as agonists. While the exo hIP_3_R3, endo hIP_3_R3 and WT HEK293 cells responded to agonist stimulation, the hIP_3_R3 E1946Q cells, like 3KO cells, did not respond to various doses of CCh (3, 10, 30 µM) (Fig. 3C, D). The percentage of responding cells were also significantly lower in stable cells with E1946Q substitution as compared to WT cells (Fig. S3A).

**Figure 3.**
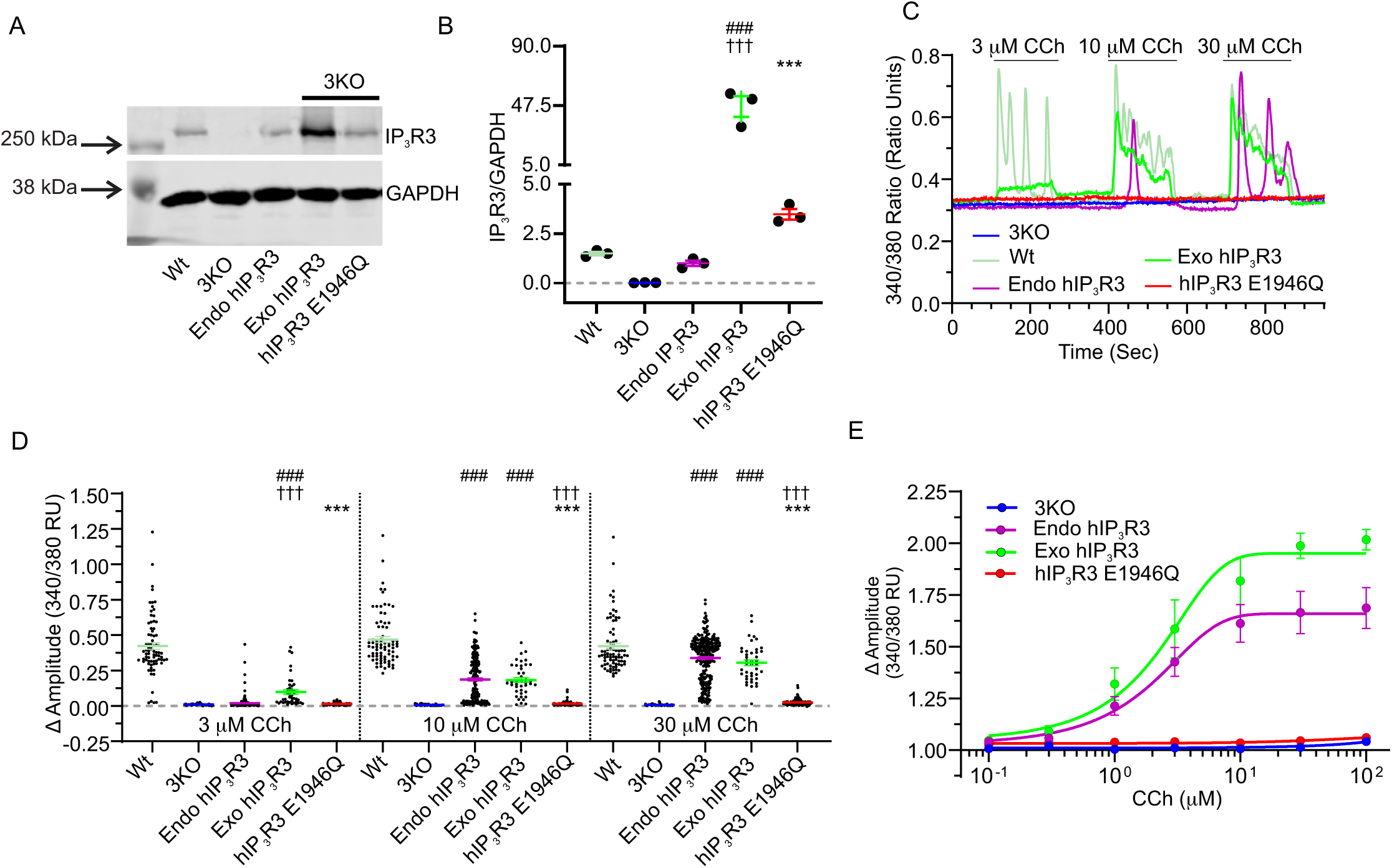
Substituting E1946 with glutamine (Q) residue in hIP_3_R3 significantly diminished agonist-induced Ca^2+^-release when stably expressed in HEK-3KO cells. Stable hIP_3_R3 cells with substitution at the E1946 position to Q were generated in HEK-3KO cells. **A**. A representative western blot depicting IP_3_R3 and GAPDH protein levels in HEK293 (WT), HEK-3KO (3KO), HEK-2KO (lacking native IP_3_R1 and IP_3_R2 sub-types, endo hIP_3_R3), 3KO cells over-expressing WT hIP_3_R3 (exo hIP_3_R3), 3KO cells over-expressing hIP_3_R3 with E1946Q substitution (hIP_3_R3 E1946Q). **B**. Scatter plot depicting quantification of IP_3_R3 protein level normalized to GAPDH from three-independent western blots. **C**. Representative traces showing CCh-induced (3, 10, 30 µM) Ca^2+^- release from 3KO (blue), Wt (pale yellow-green), endo hIP_3_R3 (purple), exo hIP_3_R3 (green) and hIP_3_R3 E1946Q (red) cells in single cell assays. **D**. Scatter plot summarizing change in amplitude (Peak ratio – Basal ratio: average of 20 ratio points immediately preceding addition) to increasing doses of CCh for experiments similar to those shown in C. **E**. Dose-response curve showing change in maximum amplitude from Fura-2/AM loaded 3KO (blue), endo hIP_3_R3 (purple), exo hIP_3_R3 (green) and hIP_3_R3 E1946Q (red) cells in population-based assays when treated with increasing concentrations (100 nM, 300 nM, 1 μM, 3 μM, 10 μM, 30 μM and 100 μM) of CCh. Data are mean ± SEM of three (N = 3) independent experiments. ###*P* < 0.001 when compared to 3KO cell line, †††*P* <0.001 when compared to endo hIP_3_R3 cell line and \*\*\**P* < 0.001 when compared to exo hIP_3_R3 cell line; one-way ANOVA with Tukey’s test was performed. Unless otherwise stated, all data above comes from at least N=3 experiments.

To further substantiate these results, we performed single cell imaging experiments using trypsin (500 nM) as an agonist. A consistent lack of trypsin-induced Ca^2+^ release in hIP_3_R3 E1946Q cells was observed when compared to WT stable cells (Fig. S3C, D). Interestingly, both the WT and endo hIP_3_R3 cells, which express comparable IP_3_R3 protein levels to hIP_3_R3 E1946Q cells, responded to trypsin (Fig. S3C, D) and various doses of CCh (Fig. 3C, D). Furthermore, we also performed population-based assays to determine CCh-induced Ca^2+^ release in hIP_3_R3 E1946Q cells. Similar to single cell imaging assays, CCh-induced Ca^2+^ release - as represented by the maximum amplitude changes (Fig. 3E) and AUC (Fig. S3B) - was abolished in hIP_3_R3 E1946Q cells when compared to both exo and endo hIP_3_R3 cells. Overall, these results indicate that the negative charge on conserved side chain residues in both the hIP_3_R1 and hIP_3_R3 sub-types is critical for coordinating Ca^2+^ and activity of hIP_3_R subtypes.

Is the negative charge on the conserved side chain residues in IP_3_R3 essential for Ca^2+^ binding and activity across species? In order to address this question, we next generated stable cell lines expressing rat IP_3_R3 (rIP_3_R3) WT or rIP_3_R3 with E1945Q substitution in HEK-3KO cells (Fig. S4A, B). We performed single cells imaging and population-based assays to determine agonist-induced Ca^2+^ release using these stable cells. Remarkably, both the CCh-(Fig. S4C-E) and trypsin-induced (Fig. S4H, I) Ca^2+^ release was decreased in rIP_3_R3 E1945Q cells as compared to rIP_3_R3 WT cells. A similar decrease in CCh-induced Ca^2+^ release was observed in population-based assays (Fig. S4F, G). To summarize, these observations indicate that the negative charge on the conserved side chain residues in IP_3_Rs across various organisms are critical for binding to Ca^2+^ and the ensuing agonist-induced Ca^2+^ release.

### Loss of negative charge in the putative Ca^2+^ binding pocket shifts the Ca^2+^ sensitivity for activation of hIP_3_R1

The previous experiments suggest strongly that negatively charged residues in a putative Ca^2+^ binding pocket centered around E2002 are necessary for the full activity of hIP_3_R, but they do not provide biophysical, mechanistic insight into how Ca^2+^ binding controls IP_3_R activity. We therefore generated stable cell lines expressing WT hIP_3_R1 or hIP_3_R1 with substitutions at the E2002 and E1938 residues in DT40-3KO chicken lymphocyte background cells (Fig. S5A, B). Interestingly, consistent with HEK-3KO cells, neutralizing the negative charge on side chain of either of these residues significantly abrogated Ca^2+^ signals evoked by trypsin in population-based assays in DT40 cells (Fig. S5C, D).

Next, we performed “on-nucleus” single channel recordings of hIP_3_R1 activity after exposure to IP_3_ and various [Ca^2+^]. As described previously, by our lab (49-51) and others (24, 52), at an optimum [ATP] (5 mM) and at a saturating [IP_3_] (10 μM), the open probability (P_o_) of the hIP_3_R increased with increasing [Ca^2+^]; channel activity was barely evident at 10 nm Ca^2+^ and increased to reach a maximum P_o_ of ∼0.7 at 200 nM-1 μM Ca^2+^. Increasing [IP_3_] to 100 μM, failed to augment activity further. At higher [Ca^2+^], hIP_3_R1 activity decreased and was essentially absent at 100 μM Ca^2+^ (Fig. 4A, F). Similar experiments were then performed in cells expressing hIP_3_R1 harboring the conservative charge mutation E2002D. While single channel activity was reduced in cells expressing this mutation (Max P_o_ ∼0.45 at 200 nM-1 μM Ca^2+^), a similar dependency on Ca^2+^ for activation and inhibition compared to WT hIP_3_R1 was retained by this mutation. The decrease in overall channel activity was not due to a reduction in the sensitivity to IP_3_, as increasing IP_3_ to 100 μM had no further effect on hIP_3_R1 P_o_ (Fig. 4B, F).

**Figure 4.**
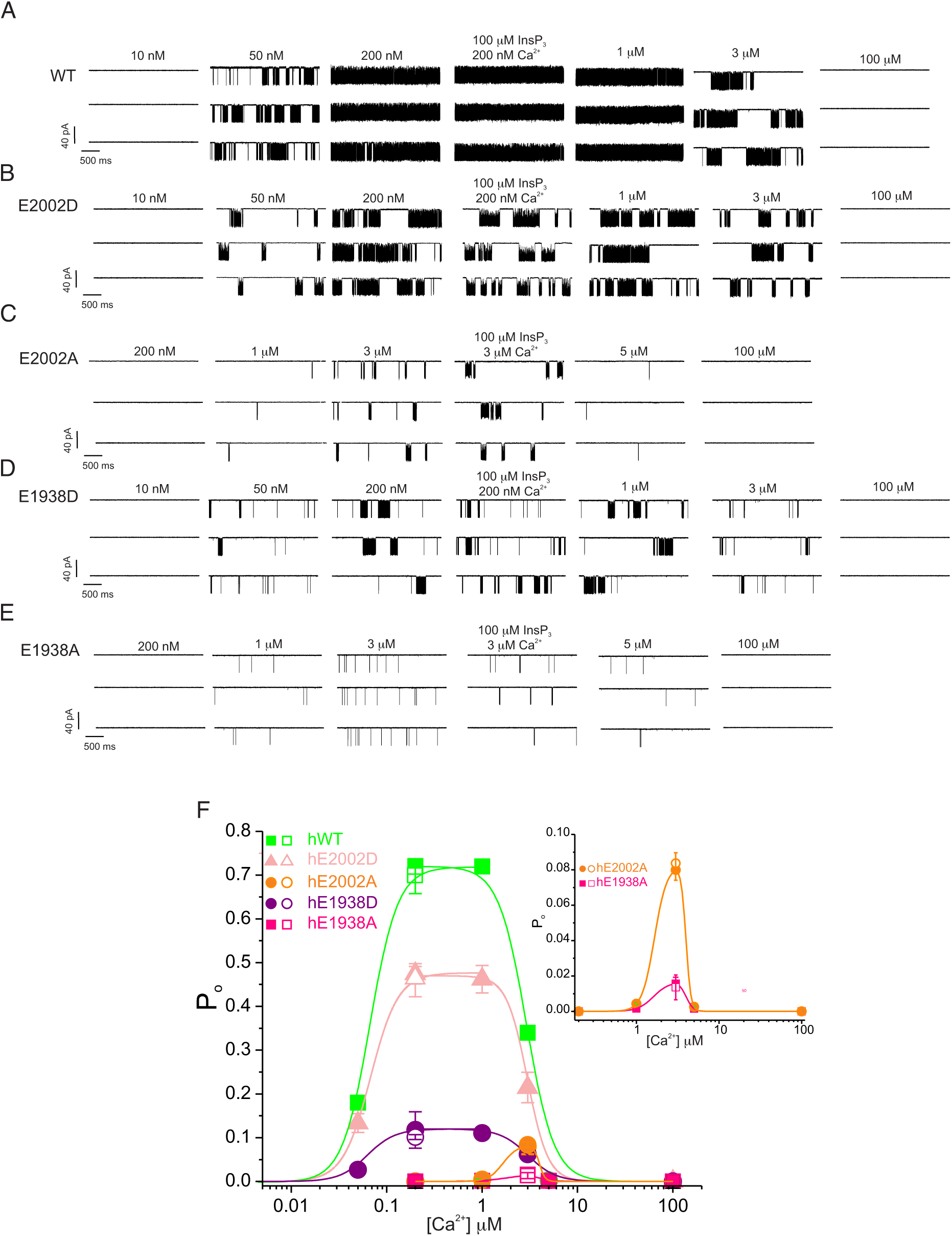
Importance of negative charges in Ca^2+^ sensor site for regulation of hIP_3_R1 by Ca^2+^. The activity of WT hIP_3_R1 and mutants was monitored in the “on-nucleus” configuration in DT40-3KO cells stably expressing the indicated constructs as detailed in methods. **A**. Representative sweeps are shown at the indicated [Ca^2+^] stimulated by 10 μM IP_3_ (unless stated otherwise) in the presence of optimal ATP for WT hIP_3_R1. Maximal activity is observed at 200 nM-1 μM Ca^2+^ and then subsequently decreases. Maximal P_o_ was not increased by increasing [IP_3_] to 100 μM. **B**. Representative activity is shown for E2002D hIP_3_R1. The maximal P_o_ at each [Ca^2+^] is reduced compared with WT hIP_3_R1 but the regulation by increasing Ca^2+^ mirrors that seen with WT hIP_3_R1. **C**. Representative activity is shown for E2002A hIP_3_R1. Maximal P_o_ is markedly reduced and the [Ca^2+^] where maximum activity is observed is right shifted to higher [Ca^2+^]. Inhibition of IP_3_R activity occurs at similar [Ca^2+^] to WT hIP_3_R1. **D**. Representative activity is shown for E1938D hIP_3_R1. In a similar fashion to E2002D, the maximal P_o_ at each [Ca^2+^] is reduced compared with WT hIP_3_R1 but the regulation by increasing Ca^2+^ mirrors that seen with WT hIP_3_R1. **E**. Representative activity is shown for E1938A hIP_3_R1. In a similar fashion to E2002A Maximal P_o_ is markedly reduced and the [Ca^2+^] where maximum activity is observed is right shifted to higher [Ca^2+^]. Inhibition of IP_3_R activity occurs at similar [Ca^2+^] to WT hIP_3_R1. **F**. Pooled data. Inset shows pooled data for alanine mutations with an expanded Y-axis.

Next, we investigated the consequences of substitution of E2002 to alanine. Strikingly, no IP_3_R activity was evident at [Ca^2+^] in a range which resulted in enhanced P_o_ in WT hIP_3_R1 (10-200 nM Ca^2+^). IP_3_R activity, albeit reduced in comparison to the hIP_3_R1 retaining negative charge at this residue, was observed at 1 μM which was further enhanced by increasing [Ca^2+^] to 3 μM and then subsequently reduced as [Ca^2+^] was increased further (Fig. 4C, F). Again, increasing the [IP_3_] to 100 μM failed to increase overall activity at optimum [Ca^2+^] for this mutation. The decreased sensitivity to activation by Ca^2+^ can be readily observed in the pooled data presented in Fig. 4F and insert. Notably, the [Ca^2+^] required for inhibition of activity was largely unaltered by this substitution. We failed to record any single channel activity when E2002 was mutated to glutamine.

As previously reported, the IP_3_R1 exhibits modal gating; increased activity is associated with increased bursting activity (49). An increase in Ca^2+^ flux is not associated with a change in open or closed times within the burst, but an increase in the duration of bursts (Fig. S6A-C). We observed that fundamental gating characteristics were not altered in any of the mutations. An increase in [Ca^2+^] both increased the burst length and concomitantly decreased the interburst interval. These data indicate that an increase in IP_3_R1 activity by Ca^2+^ is primarily associated with a destabilization of a long-closed state, while the attenuation of IP_3_R1 activity by Ca^2+^ is associated with a transition to, and stabilization of, this long closed state (Fig. S6C-G).

We also generated DT40-3KO cell lines stably expressing similar mutations at E1938 (Fig. S5A, B) which is predicted to be an additional negatively charged residue essential for coordinating Ca^2+^ in the Ca^2+^ sensing site (Fig.1). In a similar fashion to mutations in E2002, conservation of charge by substitution of E1938 for D in hIP_3_R1 resulted in a decrease in the maximum overall channel activity with no change in the absolute sensitivity to IP_3_, but the relationship for activation and inhibition of channel activity by Ca^2+^ was retained; a maximal P_o_ was attained between 200 nm-1 μM Ca^2+^ which subsequently decreased at [Ca^2+^] greater than 1 μM (Fig. 4D, F). IP_3_R1 activity in cells expressing hIP_3_R1 E1938A again displayed a right-shifted, decreased sensitivity to Ca^2+^, similar to E2002A, with maximum activity observed at 3 μM Ca^2+^ before subsequently decreasing (Fig. 4E, F). Similar to E2002D the inhibition of activity characteristic of higher [Ca^2+^] was largely unaltered when compared to WT hIP_3_R1. In total, these data are consistent with E2002 and E1938 comprising key Ca^2+^ binding residues responsible for the activation phase of the biphasic Ca^2+^ relationship underlying IP_3_R activity.

## Discussion

Regulation of IP_3_R induced Ca^2+^ release by Ca^2+^ was recognized early. Initial studies reported that IP_3_-induced Ca^2+^ release from the ER was inhibited by Ca^2+^ (53, 54). Subsequently, a series of seminal studies reported that increasing [Ca^2+^] first positively potentiates IP_3_R activity and then at higher concentrations reduces channel activity, resulting in the characteristic biphasic regulation of IP_3_-induced Ca^2+^ release and channel activity which is thought to be pivotally important for the spatio-temporal control of Ca^2+^ release (8, 20, 22, 24, 55-58). Ongoing studies have since investigated whether this fundamental regulation is directly through interaction with motifs in the IP_3_R protein itself, or through an accessory protein. For example, single channel experiments where [Ca^2+^] can be rigorously controlled on both faces of the channel, indicate that IP_3_R activity appears to be regulated by changes in [Ca^2+^]_i_ close to the open channel pore, as opposed to the [Ca^2+^] in the ER lumen (29).These data make it improbable that potential Ca^2+^ binding site/s on the luminal face of IP_3_Rs contribute to activation. Moreover, failure to detect any interactions between Ca^2+^ binding proteins and IP_3_Rs as a function of changes in the [Ca^2+^] makes it unlikely that Ca^2+^ sensitivity is conferred through an interaction with binding partner (59). Taken together these findings strongly indicate that the Ca^2+^ binding site/s lie within the receptor.

In terms of known Ca^2+^ binding motifs, IP_3_Rs lack a canonical EF-hand, C2 domain or other conventional Ca^2+^ binding motifs (27), nevertheless, Ca^2+^ binds to several linear fragments spread across various domains (35, 36). Of note, rIP_3_R1 (residues 1961-2219) and rIP_3_R2 (residues 1914-2173) coupling domain fragments expressed as recombinant proteins in bacteria strongly bound to ^45^Ca^2+^ (59). Nevertheless, residues coordinating Ca^2+^ in the IP_3_R that are responsible for modulating activity have not been functionally described. Recently, by comparing cryo-EM structures of RyR obtained in the presence and absence of its activating ligands (Ca^2+^/ATP/caffeine), putative amino acids residues forming the Ca^2+^-activating site were identified (40). Moreover, mutational analysis experimentally confirmed that these residues confer Ca^2+^ sensitivity to RyR channel (41). Cryo-EM structures of RyR resolved in presence of Ca^2+^ revealed the pore remained open, consequently, permitting the flow of Ca^2+^ ions from ER to cytoplasm (40). Using a similar approach, cryo-EM structures of hIP_3_R3 were also solved in the presence of Ca^2+^ and IP_3_. This study identified two putative cytosolic Ca^2+^ binding pockets (15) with the Ca^2+^ sensor site being highly analogous to the site validated in RyR (41, 43). Notably, the IP_3_R was reported to undergo several conformational changes in the active state as compared to its apo, unbound state. In contrast to the RyR structures, the IP_3_R3 structures were obtained at a non-physiological, high Ca^2+^ (∼2 mM) concentration and the structural changes associated with the channel pore region were not reported. We thus surmise that the channel remained closed under these conditions. As a consequence, whether either of these sites is functionally responsible for activation or inhibition of activity is unclear based on these structures (15). As noted, the residues forming the putative Ca^2+^ sensor site in hIP_3_R3 are highly conserved and similar to the Ca^2+^-activating site identified in human RyRs (Fig. 1). Moreover, Ca^2+^ binding residues at this activating site in RyR and IP_3_Rs are also highly conserved across various organisms (Fig. 1 and S1A). Even though these residues are evolutionarily well conserved, IP_3_Rs in *Capsaspora owczarzaki*, paradoxically, appears not to be regulated by Ca^2+^ in a manner similar to mammalian IP_3_R (60) and thus, this may indicate that other structural elements required for allosteric regulation of activity by Ca^2+^ are absent in this species. In the hIP_3_R3 structure, the HD/ARM2 putative Ca^2+^ binding site was poorly resolved (15). This site, while conserved in IP_3_R subtypes, is not present in RyRs (Fig. S1B). Any role for the site in regulation of IP_3_R activity should be the subject of further investigation.

In the current study, based on striking similarities between RyRs and IP_3_Rs structures at the putative Ca^2+^ sensor site, we generated several stable cell lines in HEK293-3KO and DT40-3KO cells (lacking all three native IP_3_Rs) expressing hIP_3_R1, hIP_3_R3 and rIP_3_R3 with substitutions at conserved amino acid residues forming the Ca^2+^ sensor site (Fig. 2, 3, S4, and S5). We then interrogated the channel properties of these mutants when compared to their WT counterparts. Agonist-induced Ca^2+^-release was significantly reduced when the glutamic acid residue at position 2002 in hIP_3_R1 was substituted for the native aspartate residue (Fig. 2 and S2) at this position, indicating that disruption of the structure of the putative binding pocket following reduction in size of the residue, without altering charge, disrupted Ca^2+^ regulation of IP_3_R activity. Further, agonist-induced Ca^2+^-release was completely abrogated upon neutralizing the negative charge on the glutamic acid side chain residue at the 2002 position by substitution with either glutamine or alanine (Fig. 2 and S2) indicating that the negative charge of this residue is pivotal for activity. Similar data were obtained in DT40 cells upon neutralizing negative charge on side chain residue at either E2002 or E1938 residue in hIP_3_R1 (Fig. S5).

Recent technological advances, utilizing high-speed Total Internal Reflection Microscopy, allow near electrophysiological measurement of the biophysical characteristics of Ca^2+^ release through IP_3_R clusters termed “Ca^2+^ puffs” (31, 46, 61-64). Consistent with the effect on global Ca^2+^ signals, Ca^2+^ puffs were also significantly diminished in E2002D cells or completely abrogated in E2002A and E2002Q cells (Fig. 2). Similarly, substitution of glutamine for glutamic acid at the analogous 1946 position in hIP_3_R3 also decimated agonist-induced Ca^2+^-release (Fig. 3 and S3). A similar result was obtained mutating the analogous residue in rIP_3_R3 (Fig. S4). We did not consider substitution of multiple Ca^2+^ coordinating residues in a site to avoid excessive perturbations in the native structure of IP_3_Rs.

Our single channel data provides unparalleled mechanistic insight into how the binding of Ca^2+^ to the Ca^2+^ sensor site influences the activity of the IP_3_R. First, mutation of either E2002 or E1938 by substitution with an aspartate acid residue (Fig. 4B, D) or alanine residue (Fig. 4C, E) did not alter the single channel conductance or the modal gating of the channel in bursts, suggesting that these mutations did not grossly alter the structure of the hIP_3_R1, or the conformational changes required from IP_3_ binding to opening of the channel. Notably, however, mutation of these residues with either substitution, had marked effects on channel P_o_ and most strikingly the loss of negative charge markedly altered the activation of channel activity by Ca^2+^ without altering the inhibition at higher [Ca^2+^] (Fig. 4A-E). This is most clearly appreciated by comparing the pooled activity relationships for WT hIP_3_R1 with the charge conserved aspartic acid substitutions at either site with the mutations to alanine (Fig. 4F). Clearly, while the activity of the aspartic acid (Fig. 4B, D) substituted mutants are reduced compared to WT hIP_3_R1 (Fig. 4A), the activity is regulated in an identical biphasic manner. In contrast, the alanine mutants have further reduced maximum activity, but peak activation is achieved at higher [Ca^2+^] when compared to WT or glutamic acid harboring mutants (Fig. 4C, E). The reduction in overall activity of each mutant, compared to WT is consistent with disruption of the overall integrity of the Ca^2+^ binding pocket by alteration in the size and charge of side chains resulting in disordered transduction of conformational changes on Ca^2+^ binding to gating of the pore. Primarily, these data, further reinforce that the negative charge on both amino acid residues E2002 and E1938 are critically important for coordinating Ca^2+^ in the Ca^2+^ sensor site which leads to the activation of IP_3_R. Further work is necessary to elucidate the structural basis of inhibition of IP_3_R activity by Ca^2+^.

Limitations of our study: Each protomer contributes one Ca^2+^ binding site at the ARM3/LNK interface to the tetrameric assembly and Ca^2+^ regulation of IP_3_R activity is positively cooperative, thus whether Ca^2+^ binding to all four Ca^2+^ sensor site in homo-/hetero-tetramer is absolutely required for pore opening/channel activation is yet to be determined. Additionally, as IP_3_R sub-types have differential affinity for IP_3_ and ATP, it is yet to be determined whether the affinities of Ca^2+^ binding sites are different for individual subtypes and how Ca^2+^ shapes agonist-induced Ca^2+^ release from blended IP_3_R sub-types (hetero-tetramers).

## Methods and Materials

### Plasmid constructs

Substitutions to amino acids aspartate (D), alanine (A) or glutamine (Q) at the desired site/residue in rIP_3_R3, hIP_3_R3 and hIP_3_R1 were generated using *Pfu* Ultra II Hotstart 2X Master Mix (Agilent Technologies) and appropriate primers obtained from Integrated DNA Technologies (Table. S1). A Quikchange lighting site-directed mutagenesis kit (Agilent Technologies #210518) was used to introduce the desired substitution in cDNAs encoding the rIP_3_R3 (NP_037270), hIP_3_R3 (NP_002215.2) and hIP_3_R1 (NP_001093422.2) in pDNA3.1 expression plasmid using mutagenic primers. The introduction of desired substitution and coding regions for all the constructs were confirmed by Sanger sequencing.

### Alignment of IP_3_R protein sequences from various organisms

Alignment of IP_3_R protein sequences from: *Rattus norvigecus* - short IP_3_R1 (used in Fig.1E) (P29994.2), IP_3_R1 (NP_001257525.1) (used in Fig.S1), IP_3_R2 (NP_112308.1) and IP_3_R3 (NP_037270.2); *Homo sapiens* - IP_3_R1 (NP_001093422.2), IP_3_R2 (NP_002214.2), IP_3_R3 (NP_002215.2), RyR1 (NP_000531.2), RyR2 (NP_001026.2), RyR3 (NP_001027.3); *Capsaspora owczarzaki* - IP_3_RA (XP_004347577.1); *Caenorhabditis elegans* - IP_3_R1 (NP_001023170.1); *Drosophila melanogaster* - IP_3_R (NP_730941.1) ; *Danio rerio* - IP_3_R1 (XP_021335554.1); *Gallus gallus* - IP_3_R1 (NP_001167530.1); *Mus musculus* - IP_3_R1 (NP_034715.3); *Bos taurus* - IP_3_R1 (NP_777266.1); *Canis familiaris* - IP_3_R1 (XP_005632286.1); *Macaca mulatta* - IP_3_R1 (NP_034715.3); and *Pan troglodytes* - IP_3_R1 (XP_009443057.1) were generated using GeneDoc.

### Cell culture, transfection and generation of stable cell lines

DT40-3KO, chicken B lymphocyte cells engineered through homologous recombination for the deletion of all the three-native endogenous IP_3_R isoforms (65), were cultured in RPMI 1640 media supplemented with 1% chicken serum, 10% fetal bovine serum, 100 U/ml penicillin, 100 μg/ml streptomycin in an incubator set to 39°C with 5% CO_2_. Transfections in DT40-3KO were performed as previously described (66). In brief, 5 million cells were washed with phosphate buffered saline (PBS) and electroporated with 4-6 μg of appropriate plasmid construct using an Amaxa cell nucleofector (Lonza laboratories) and nucleofection reagent (362.88 mM ATP-disodium salt, 590.26 mM MgCl_2_ 6.H_2_O, 146.97 mM KH_2_PO_4_, 23.81mM NaHCO_3_, and 3.7 mM glucose at pH 7.4). Following transfection, the cells were allowed to recover for 24 h and subsequently, transferred into 96 well-plates containing media supplemented with 2 mg/ml G418. Next, 10-14 days after transfection, clones expressing the desired construct were expanded and subsequently screened by western blotting. Cell lines stably expressing the construct were used in further experiments.

HEK-3KO, HEK293 cells engineered in our laboratory using CRISPR/Cas technology for the deletion of all the three-native endogenous IP_3_R isoforms (19) were cultured in DMEM media supplemented with 10% fetal bovine serum, 100 U/ml penicillin, 100 μg/ml streptomycin in an incubator set to 37°C with 5% CO_2_. Transfections of the appropriate plasmid construct was performed using a protocol previously described (19). In brief, 1 million cells were washed with PBS and electroporated with 5-10 μg of appropriate plasmid using Amaxa cell nucleofector kit T (Lonza laboratories). Cells were allowed to recover for 48 h, and subsequently, sub cultured into new 10 cm^2^ plates containing media supplemented with 1.5-2 mg/ml G418. Following 7 days of selection, individual colonies of cells were picked and transferred to new 24-well plates containing media supplemented with 1.5-2 mg/ml G418. Clonal lines were expanded and those expressing the desired constructs were confirmed by western blotting (18).

### Western blotting

For western blotting, total protein was isolated from indicated control and stable cell lines using membrane-bound extraction buffer (10 mM Tris-HCl, 10 mM NaCl, 1 mM EGTA, 1 mM EDTA, 1 mM NaF, 20 mM Na_4_P_2_O_7_, 2 mM Na_3_VO_4_, 1% Triton X-100 (v/v), 0.5% sodium deoxycholate (w/v), and 10% glycerol) supplemented with protease inhibitors (Roche, USA). Briefly for protein isolation, following addition of appropriate amount of lysis buffer, cells were harvested in 1.5 ml tubes and placed on ice for 30 minutes. In order to disrupt the pellet, the tubes were vortexed for 10 seconds every 10 minutes and returned on ice. Following incubation on ice, the cell lysates were centrifuged at 13,000 rpm at 4°C for 10 minutes. The supernatant was transferred to new labeled tubes. Protein concentration in the lysates was estimated using the D_c_ protein assay kit (Bio-Rad). Equal amounts of lysates were then subjected to SDS-PAGE and transferred to a nitrocellulose membrane. The membranes were incubated with indicated primary antibodies and appropriate secondary antibodies before imaging with an Odyssey infrared imaging system (LICOR Biosciences). Band intensities were quantified using Image Studio Lite Ver 5.2 and presented as ratios of IP_3_R to GAPDH. The IP_3_R1 antibody (#ARC154, Antibody Research Corporation) was used at 1:1000 dilution, IP_3_R3 antibody (#610313, BD Transduction Laboratories) was used at 1:1000 dilution, GAPDH (#AM4300, Invitrogen) was used at 1:75,000 dilution, secondary goat anti-rabbit (SA535571, Invitrogen) and secondary goat anti-mouse (SA535521, Invitrogen) antibodies were used at 1:10,000 dilution (63).

### Measurement of cytosolic Ca^2+^ in intact cells

Population-based Ca^2+^ imaging in the indicated cell lines was performed as described previously (18, 63). Briefly, adherent cells were plated in 10 cm^2^ cell culture dishes. Upon attaining 90-100% confluency, the cells were loaded with 4 μM Fura-2/AM in cell culture media and incubated at 37°C in the dark for an hour. The cells were subsequently washed three times with imaging buffer (10 mM HEPES, 1.26 mM Ca^2+^, 137 mM NaCl, 4.7 mM KCl, 5.5 mM glucose, 1 mM Na_2_HPO_4_, 0.56 mM MgCl_2_, at pH 7.4). An equal number (300,000 cells/well) of cells were seeded into each well of a black-walled 96-well plate. In contrast, suspended cells were cultured in 75 cm^2^ flasks, pelleted and washed once with imaging buffer prior to counting. An equal number (500,000 cells/well) of cells were incubated with 4 μM Fura-2/AM in imaging buffer at room temperature for one hour with constant rocking. Subsequently, the cells were pelleted and washed twice prior to being seeded into wells of a black-walled 96-well plate. The cells were centrifuged at 200g for 2 minutes to plate the bottom of the wells and incubated at appropriate temperature for 30 minutes prior to imaging. Fura-2/AM imaging was carried out by alternatively exciting the loaded cells between 340 nm and 380 nm; emission was monitored at 510 nm using FlexStation 3 (Molecular Devices). Data was exported to Microsoft Excel where peak response to increasing concentrations of agonist (0.1–100 μM CCh or 0.01–3 μM Trypsin) was determined by calculating the 340/380 ratio and normalizing it to the average of the first 5 data points of the experiment (the basal Ca^2+^ value). Area under the curve was calculated in GraphPad Prism 8 as increases above baseline that are greater than 10% of the distance from the minimum to the maximum Y using the five data points prior to agonist addition as the baseline. Curve fitting was performed using a logistic dose-response equation in GraphPad Prism 8. Data reported as triplicates from at least three individual plates.

Single cell Ca^2+^ imaging in indicated cell lines was performed as described previously. Briefly, cells were seeded on 15-mm glass coverslips in 12 well-plates and left undisturbed overnight. Once attached to the coverslip, the cells were washed with imaging buffer prior to attachment of the glass coverslip to a Warner perfusion chamber using vacuum grease. Subsequently, the cells were loaded with 2 μM Fura-2/AM for 25 minutes in the dark at room temperature. Cells were then perfused with imaging buffer and stimulated with indicated concentrations of CCh or Trypsin. Ca^2+^ imaging was performed using an inverted epifluorescence Nikon microscope equipped with a 40-x oil immersion objective. Fura-2/AM imaging was carried out by alternatively exciting the loaded cells between 340 nm and 380 nm; emission was monitored at 505 nm. Images were captured every second with an exposure of 20 ms and 4 × 4 binning using a digital camera driven by TILL Photonics software as previously described (18). Image acquisition was performed using TILLvisION software and data was exported to Microsoft Excel where data was analyzed for change in peak amplitude and percentage of cells with pre-defined peak amplitudes. Each experiment was repeated at least three times.

### Detection and analysis of Ca^2+^ puffs using TIRF microscopy

Stable hIP_3_R1 exogenous WT cells or cells with substitutions at the E2002 site were cultured on 15-mm glass coverslips coated with poly-D-lysine (100 μg/ml) in a 35-mm dish for 36 hours. Prior to imaging, the cells were washed three times with imaging buffer. The cells were subsequently incubated with Cal520-AM (5 µM; AAT Bioquest #21130) and ci-IP_3_/PM (1 μM, Tocris #6210) in imaging buffer with 0.01 % BSA in dark at room temperature. After 1-hour incubation, the cells were washed three times with imaging buffer and incubated in imaging buffer containing EGTA-AM (5 μM, Invitrogen #E1219). After 45 minutes incubation, the media was replaced with fresh imaging buffer and incubated for additional 30 minutes at room temperature to allow for de-esterification of loaded reagents.

Following loading, the coverslip was mounted in a chamber and imaged using an Olympus IX81 inverted Total Internal Reflection Fluorescence Microscopy (TIRFM) equipped with oil-immersion PLAPO OTIRFM 60x objective lens/1.45 numerical aperture. Olympus CellSens Dimensions 2.3 (Build 189987) software was used for imaging. The cells were illuminated using a 488 nm laser to excite Cal-520 and the emitted fluorescence was collected through a band-pass filter by a Hamamatsu ORCA-Fusion CMOS camera. The angle of the excitation beam was adjusted to achieve TIRF with a penetration depth of ∼140 nm. Images were captured from a final field of 86.7 µm x 86.7 µm (400 × 400 pixels, one pixel=216 nm) at a rate of ∼50 frames/second (binning 2 × 2) by directly streaming into RAM. To photo-release IP_3_, UV light from a laser was introduced to uniformly illuminate the field of view. Both the intensity of the UV flash and the duration (1 second) for uncaging IP_3_ were optimized to prevent spontaneous puffs in the absence of loaded ci-IP_3_. Images were exported as vsi files. Images, 5 seconds prior and 60 seconds after flash photolysis of ci-IP_3_, were captured, as described previously (63).

The vsi files were converted to tiff files using Fiji and further processed using FLIKA, a Python programming-based tool for image processing (67). From each recording, ∼100 frames (∼2 seconds) before photolysis of ci-IP_3_ were averaged to obtain a ratio image stack (F/Fo) and standard deviation for each pixel for recording up to 13 seconds following photolysis. The image stack was Gaussian-filtered, and pixels that exceeded a critical value (1.0 for our analysis) were located. The ‘Detect-puffs’ plug-in was utilized to detect the number of clusters (puff sites), number of events (number of puffs), amplitudes and durations of localized Ca^2+^ signals from individual cells. All the puffs identified automatically by the algorithm were manually confirmed before analysis. The results from FLIKA were saved to Microsoft Excel and graphs were plotted using GraphPad Prism 8 (68).

### Preparation of DT40 cell nuclei

Isolated DT40 nuclei were prepared using homogenization. The homogenization buffer (HB) contained 250 mM sucrose, 150 mM KCl, 3 mM 2-mercaptoethanol (β-ME), 10 mM Tris, 1 mM Phenylmethanesulphonylfluoride (PMSF), pH 7.5 with a complete protease inhibitor tablet (Roche, USA). Cells were washed and resuspended in HB prior to nuclear isolation using an RZR 2021 homogenizer (Heidolph Instruments, Germany) with 15 strokes at 1200 rpm. A 3 μl aliquot of nuclear suspension was placed in 3 ml of bath solution (BS) which contained 140 mM KCl, 10 mM HEPES, 500 μM BAPTA and 246 nM free Ca^2+^, pH 7.1. Nuclei were allowed to adhere to a plastic culture dish for 10 minutes prior to patching.

### On-nuclei patch clamp experiments

Single IP_3_R channel potassium currents (i_k_) were measured in the on-nucleus patch clamp configuration using pCLAMP 9 and an Axopatch 200B amplifier (Molecular Devices, Sunnydale, CA, USA). Pipette solution contained 140 mM KCl, 10 mM HEPES, 5 mM ATP, with varying concentrations of IP_3_, BAPTA and free Ca^2+^. Free [Ca^2+^] was calculated using Max Chelator freeware and verified fluorometrically. Traces were consecutive 3 second sweeps recorded at −100 mV, sampled at 20 kHz and filtered at 5 kHz. A minimum of 15 seconds of recordings were considered for data analyses. The data are representative of between 3 and 5 experiments for each condition presented. Pipette resistances were typically 20 MΩ and seal resistances were > 5 GΩ (49).

## Data analysis

Single channel openings were detected by half-threshold crossing criteria using the event detection protocol in Clampfit 9. We assumed that the number of channels in any particular nuclear patch is represented by the maximum number of discrete stacked events observed during the experiment. Even at low P_o_, stacking events were evident. Only patches with one apparent channel were considered for analyses. Probability of opening (P_o_), unitary current (i_k_), open and closed times, and burst analyses were calculated using Clampfit 9 and Origin 6 software (Origin Lab, Northampton, MA, USA). All-points current amplitude histograms were generated from the current records and fitted with a normal Gaussian probability distribution function. The coefficient of determination (R^2^) for every fit was *>*0.95. The P_o_ was calculated using the multimodal distribution for the open and closed current levels. The threshold for an open event was set at 50% of the maximum open current and events shorter than 0.1 ms were ignored. A ‘burst’ was defined as a period of channel opening following a period of no channel activity which was greater than five times the mean closed time (0.2 ms) within a burst. Ca^2+^ dependency curves were fitted separately for activation and inhibition with the logistic equation:

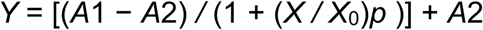

where *A*1 and *A*2 are asymptotes, *X* is the concentration of Ca^2+^, *X*_0_ is the half-maximal concentration and *p* is the slope related to the Hill coefficient. Equation parameters were estimated using a non-linear, least-squares algorithm. Two-tailed heteroscedastic t*-*tests with P values < 0.05 were considered to have statistical significance.

## Statistical Analysis

All statistical tests were conducted in GraphPad Prism 8 and data are presented as the mean ± SEM. Statistical significance was determined using one-way ANOVA with Tukey’s test.

## Acknowledgements

The authors wish to thank Dr. Kamil J Alzayady, Dr. Sundeep Malik, Ms. Taylor R. Knebel for generating stable cell lines and all the members of Yule lab especially Ms. Kai-Ting Huang and Ms. Amanda M Wahl for their valuable suggestions. The authors wish to thank Dr. Ian Parker (UC Irvine) and Dr. Jeffrey Lock (UC Irvine) for assistance and advice with FLIKA.

## Competing interests

The authors of this work have no competing interests to disclose.

## Funding

This work was supported by National Institutes of Health Grant NIH/DE019245 to Dr. David I. Yule.

## Supplementary figures

**Figure S1.**
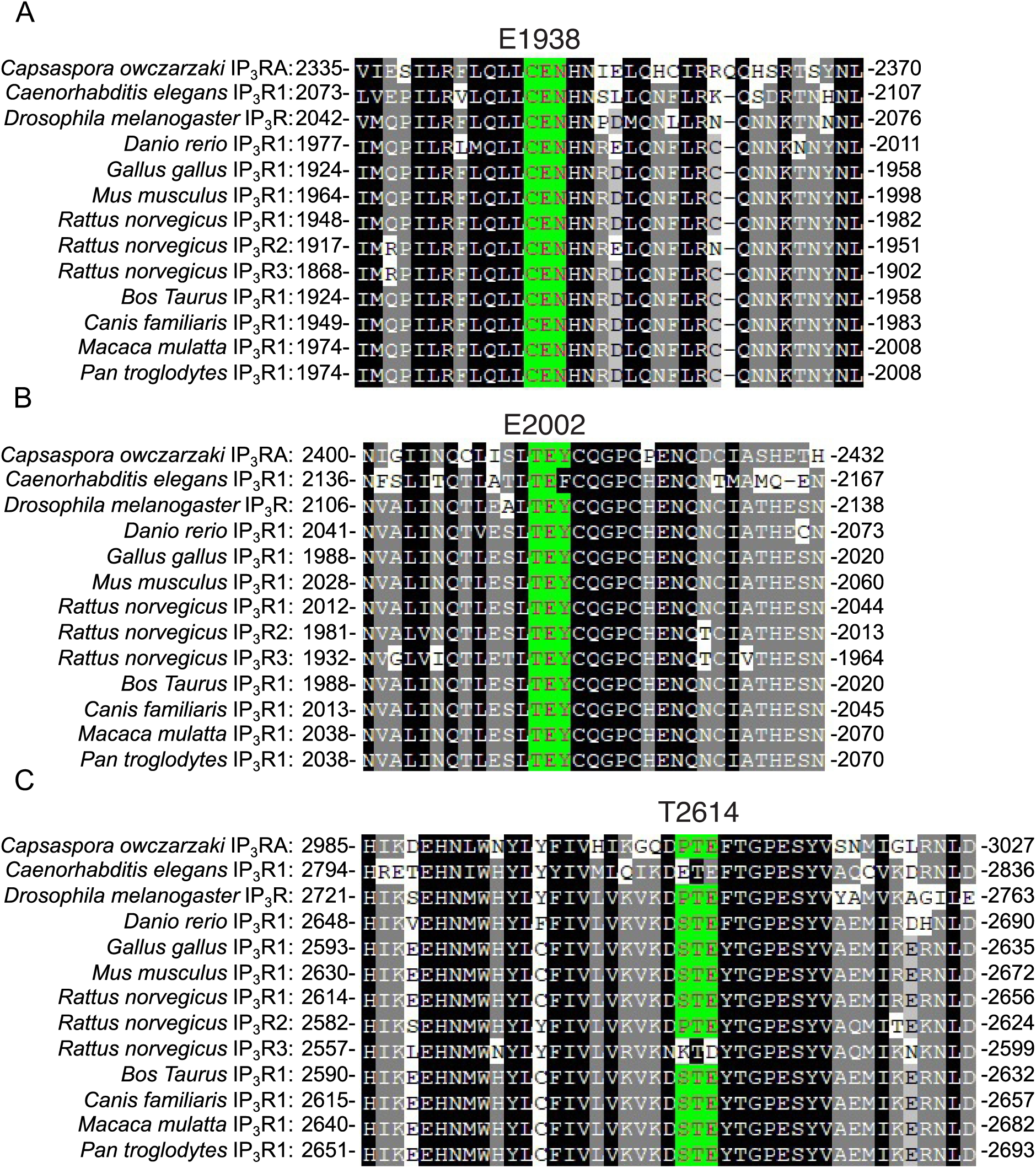
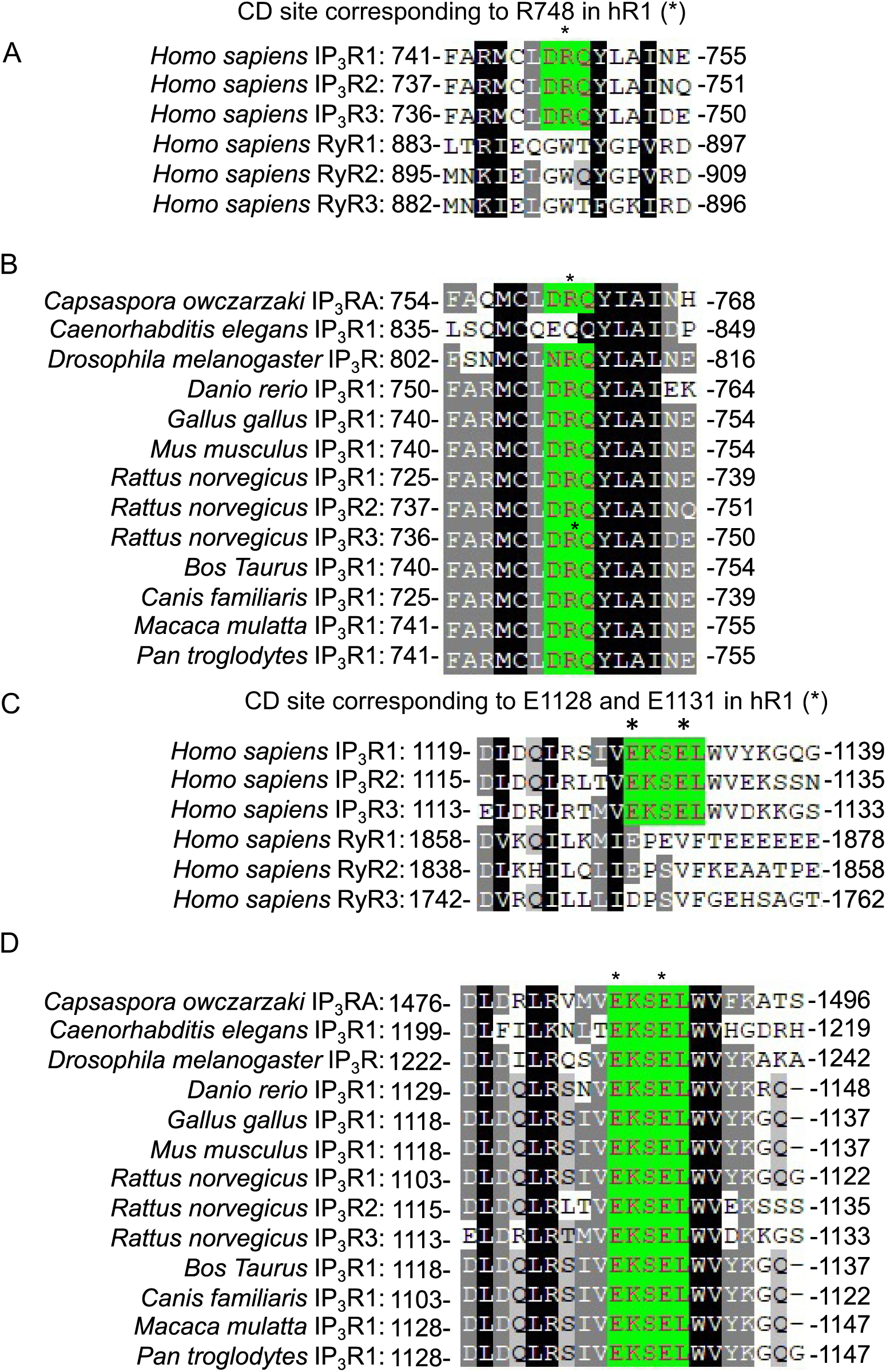
**A**. Putative Ca^2+^ coordinating residues in hIP_3_Rs are highly conserved across various organisms. The E1938, E2002 and T2614 residues (highlighted in green and indicated by asterisks) in hIP_3_R1 are conserved across various organisms. **S1. B**. The R748, E1128 and T1131 residues (highlighted in green and indicated by asterisks) in hIP_3_R1 are conserved across various organisms, but not in hRyRs.

**Figure S2.**
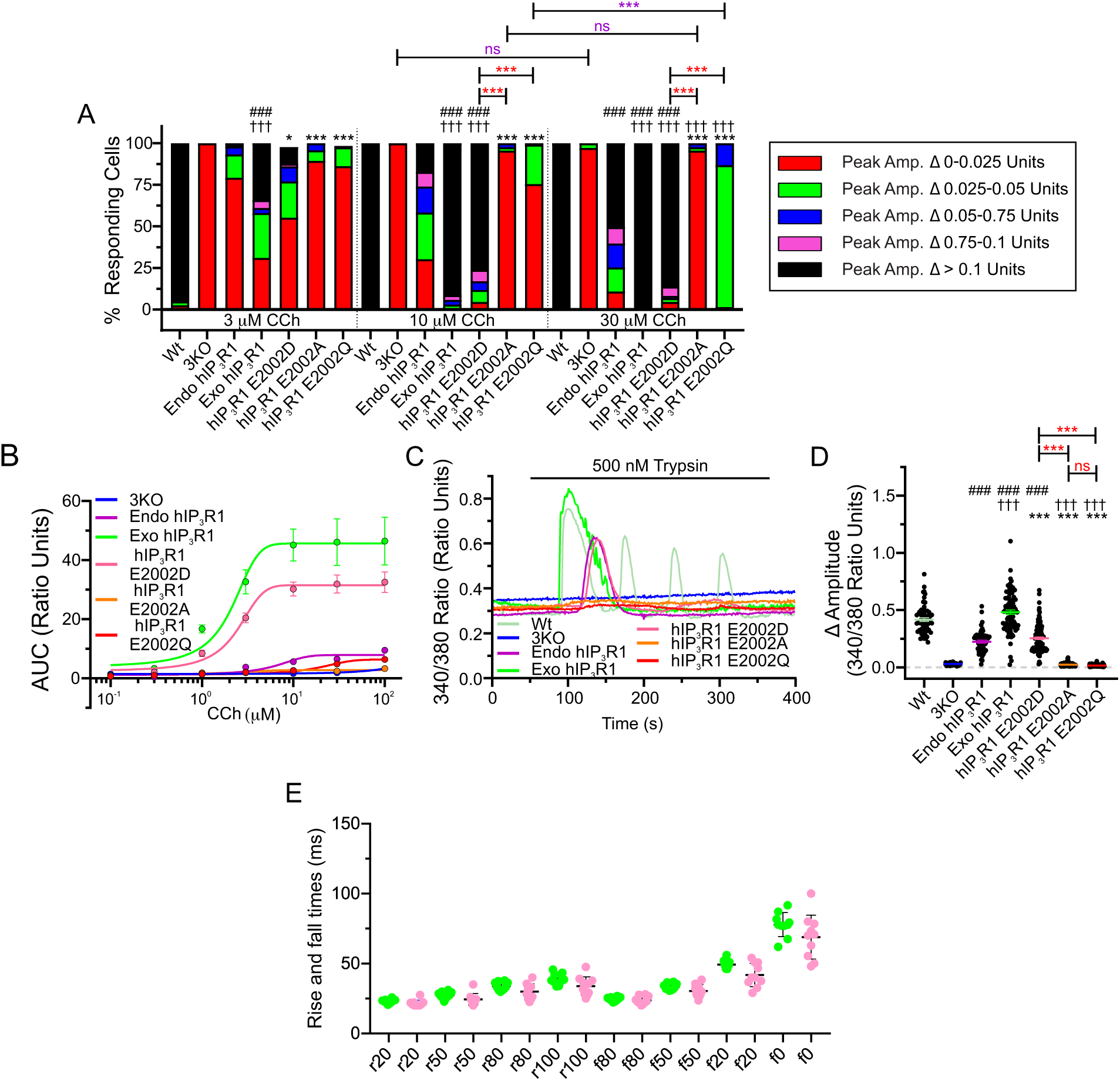
**A**. Stacked bar graph summarizing changes in amplitudes from WT HEK293, 3KO, indicated hIP_3_R1 stable cell lines in single cell assays segregated into pre-determined ranges upon stimulation with (3 μM, 10 μM and 30 μM) CCh. Cells with an amplitude change greater than 0.1 ratio units (black portion of bars) are considered as cells respondering to CCh stimulation. **B**. Dose-response curves depicting AUC from Fura-2/AM loaded 3KO (blue), endo hIP_3_R1 (purple), exo hIP_3_R1 (green), hIP_3_R1 E2002D (pink), hIP_3_R1 E2002A (orange) and hIP_3_R1 E2002Q (red) cells in population-based assays when treated with increasing concentrations (100 nM, 300 nM, 1 μM, 3 μM, 10 μM, 30 μM and 100 μM) of CCh. **C**. Representative traces showing trypsin-induced (500 nM) Ca^2+^-release from 3KO (blue), Wt (pale yellow-green), endo hIP_3_R1 (purple), exo hIP_3_R1 (green), hIP_3_R1 E2002D (pink), hIP_3_R1 E2002A (orange), hIP_3_R1 E2002Q (red) cells in single cell assays. **D**. Scatter plot summarizing change in amplitude (Peak ratio – Basal ratio: average of 20 ratio points immediately preceding addition) upon stimulation with trypsin (500 nM) for experiments similar to those shown in C. **E**. The mean time of a 20%, 50%, 80%, or 100% rise (r) and decay (f) regarding the fluorescence of Ca^2+^ puffs (n=10 cells) from exo hIP_3_R1 cells (green), hIP_3_R1 E2002D (pink). Data are mean ± SEM of three (N = 3) independent experiments. ###*P* < 0.001 when compared to 3KO cell line, †††*P* < 0.001 when compared to endo hIP_3_R1 cell line and \**P* < 0.05, \*\*\**P* < 0.001 when compared to exo hIP_3_R1 cell line; one-way ANOVA with Tukey’s test was performed. ns; not significant. Unless otherwise stated, all data above comes from at least N=3 experiments.

**Figure S3.**
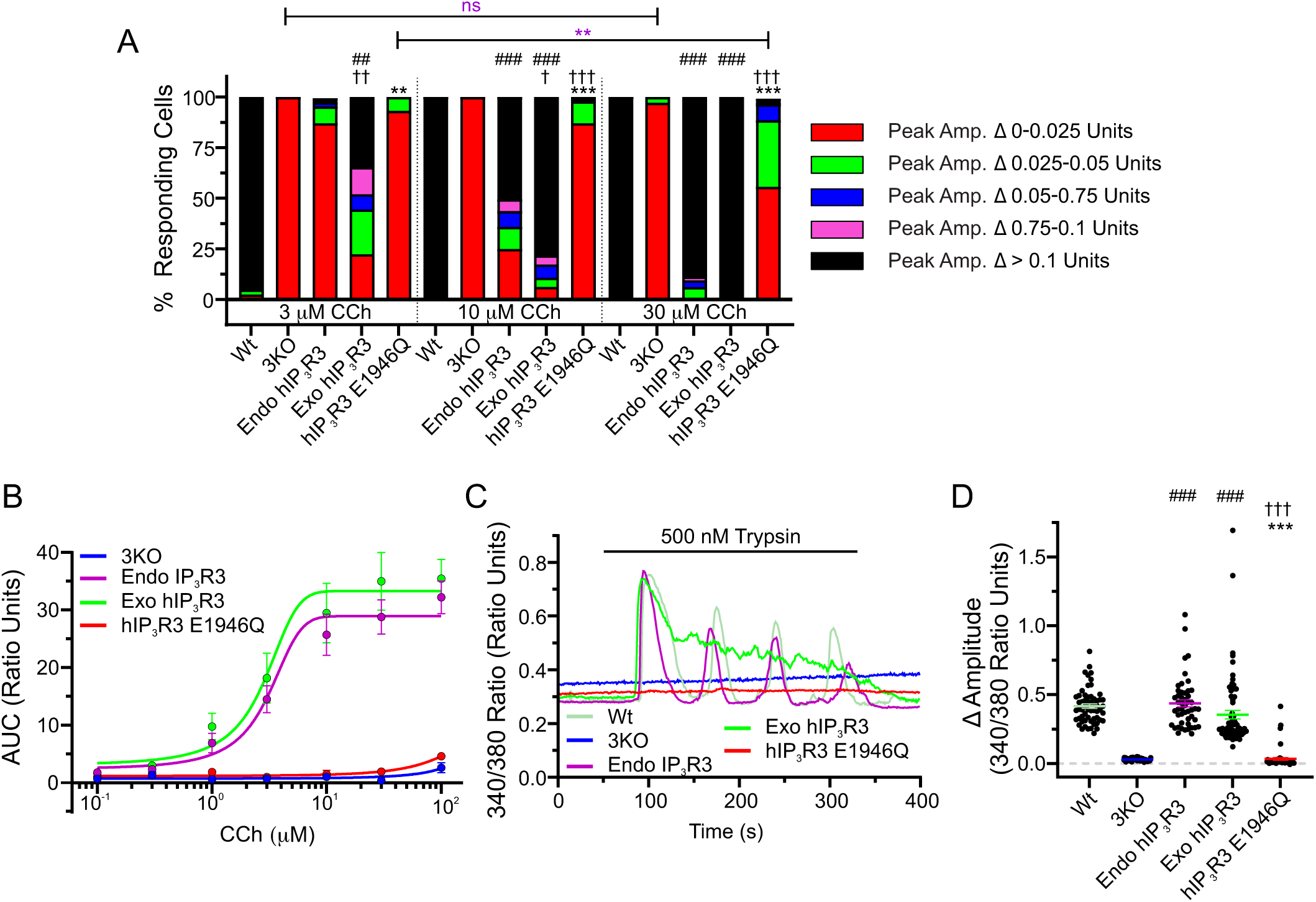
**A**. Stacked bar graph summarizing changes in amplitudes from WT HEK293, 3KO, indicated hIP_3_R3 stable cell lines in single cell assays segregated into pre-determined ranges upon stimulation with (3 μM, 10 μM and 30 μM) CCh. Cells with an amplitude change greater than 0.1 ratio units (black portion of bars) are considered as cells responding to CCh stimulation. **B**. Dose-response curves depicting AUC from Fura-2/AM loaded 3KO (blue), endo hIP_3_R3 (purple), exo hIP_3_R3 (green) and hIP_3_R3 E1946Q (red) cells in population-based assays when treated with increasing concentrations (100 nM, 300 nM, 1 μM, 3 μM, 10 μM, 30 μM and 100 μM) of CCh. **C**. Representative traces showing trypsin-induced (500 nM) Ca^2+^-release from 3KO (blue), Wt (pale yellow-green), endo hIP_3_R3 (purple), exo hIP_3_R3 (green) and hIP_3_R3 E1946Q (red) cells in single cell assays. **D**. Scatter plot summarizing change in amplitude (Peak ratio – Basal ratio: average of 20 ratio points immediately preceding addition) upon stimulation with trypsin (500 nM) for experiments similar to those shown in C. Data are mean ± SEM of three (N = 3) independent experiments. ##*P* < 0.01, ###*P* < 0.001 when compared to 3KO cell line, †*P* < 0.05, ††*P* < 0.01, †††*P* < 0.001 when compared to endo hIP_3_R1 cell line, \*\**P* < 0.01, \*\*\**P* < 0.001 when compared to exo hIP_3_R3 cell line and \*\**P* (purple) < 0.01 when percentage of cells with change in amplitude > 0.025 are compared to same cell line at different concentrations of agonist; one-way ANOVA with Tukey’s test was performed. ns; not significant. Unless otherwise stated, all data above comes from at least N=3 experiments.

**Figure S4.**
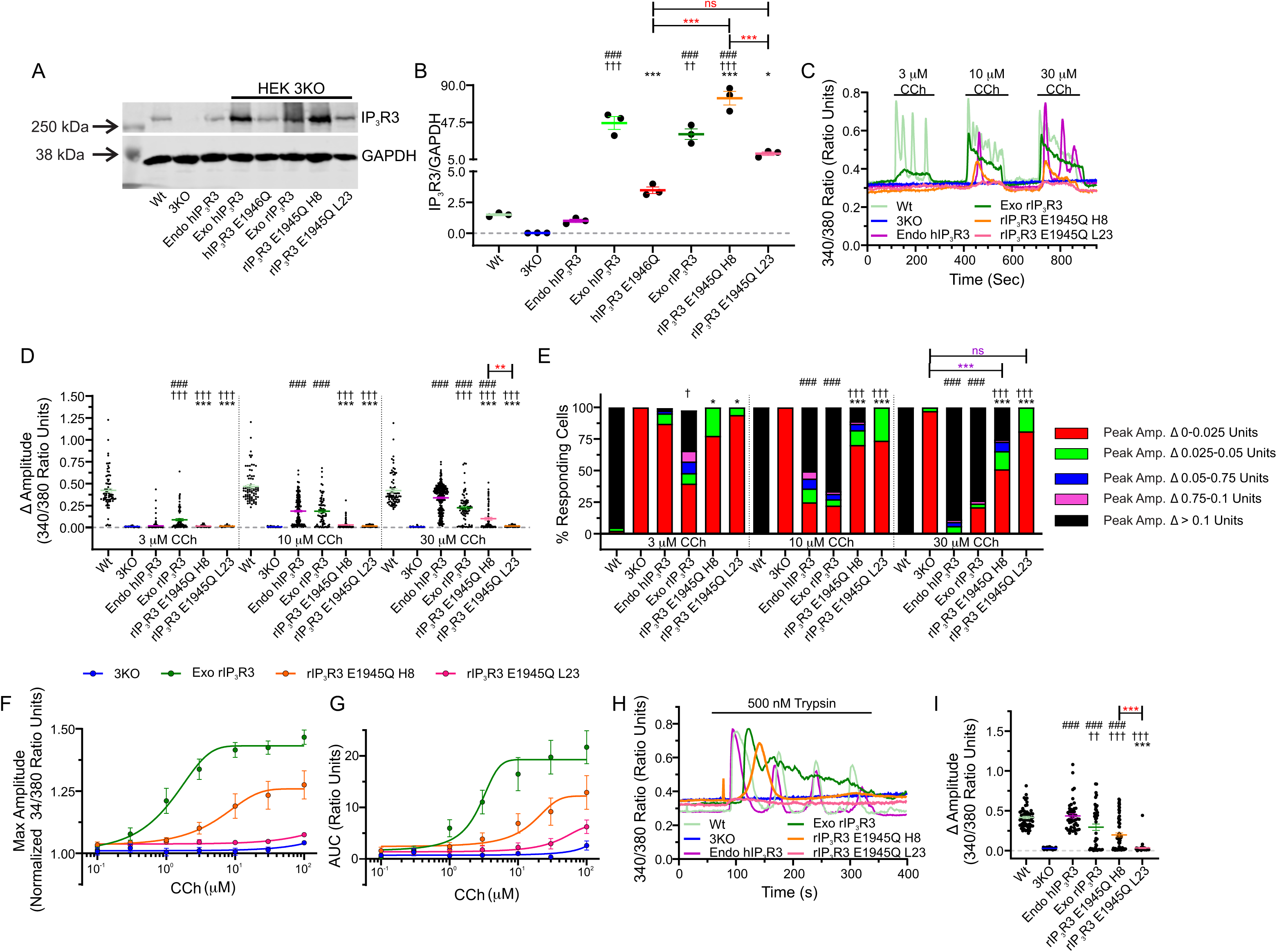
Substituting E1945 residue with glutamine (Q) in rIP_3_R3 significantly diminished agonist-induced Ca^2+^-release when stably expressed in HEK-3KO cells. Stable rat IP_3_R3 cell lines with substitution at the E1945 position to Q were generated in HEK-3KO cells. **A**. Western blot depicting IP_3_R protein levels in HEK-3KO cells over-expressing WT rIP_3_R3 (exo rIP_3_R3) and rIP_3_R3 with E1945Q substitution (rIP_3_R3 E1945Q H8 and L23). **B**. Scatter plot depicting quantification of IP_3_R3 protein levels from three-independent western blots. **C**. Representative traces showing CCh-induced (3, 10, 30 µM) Ca^2+^-release from 3KO (blue), Wt (pale yellow-green), endo hIP_3_R3 (purple), exo rIP_3_R3 (green), rIP_3_R3 E1945Q (pink, orange) cells in single cell assays. **D**. Scatter plot summarizing change in amplitude (Peak ratio – Basal ratio: average of 20 ratio points immediately preceding addition) to increasing doses of CCh for experiments similar to those shown in C. **E**. Stacked bar graph summarizing changes in amplitudes from WT HEK293, 3KO, endo hIP_3_R3 and indicated rIP_3_R3 stable cell lines in single cell assays segregated into pre-determined ranges upon stimulation with (3 μM, 10 μM and 30 μM) CCh. Cells with an amplitude change greater than 0.1 ratio units (black portion of bars) are considered as responders to CCh stimulation. Dose-response curve showing **F**. maximum amplitude and **G**. AUC from Fura-2/AM loaded 3KO (blue), exo rIP_3_R3 (green) and rIP_3_R3 E1945Q (orange, pink) cells in population-based assays when treated with increasing concentrations (100 nM, 300 nM, 1 μM, 3 μM, 10 μM, 30 μM and 100 μM) of CCh. **H**. Representative traces showing trypsin-induced (500 nM) Ca^2+^ release from 3KO (blue), Wt (pale yellow-green), endo hIP_3_R3 (purple), exo rIP_3_R3 (green) and rIP_3_R3 E1945Q (orange, pink) cells in single cell assays. **I**. Scatter plot summarizing change in amplitude (Peak ratio – Basal ratio: average of 20 ratio points immediately preceding addition) upon stimulation with trypsin (500 nM) for experiments similar to those shown in H. Data are mean ± SEM of three (N = 3) independent experiments., ###*P* < 0.001 when compared to 3KO cell line and †*P* < 0.05, ††P < 0.01, †††*P* <0.001 when compared to endo hIP_3_R3 cell line, \**P* < 0.05, \*\**P* < 0.01, \*\*\**P* < 0.001 when compared to exo rIP_3_R3 cell line, \*\**P* (red) < 0.01, \*\*\**P* (red) < 0.001 when compared to other mutant cell lines and \*\*\**P* (purple) < 0.001 when percentage of cells with change in amplitude > 0.025 are compared to same cell line at different concentrations of agonist; one-way ANOVA with Tukey’s test was performed. Unless otherwise stated, all data above comes from at least N=3 experiments.

**Figure S5.**
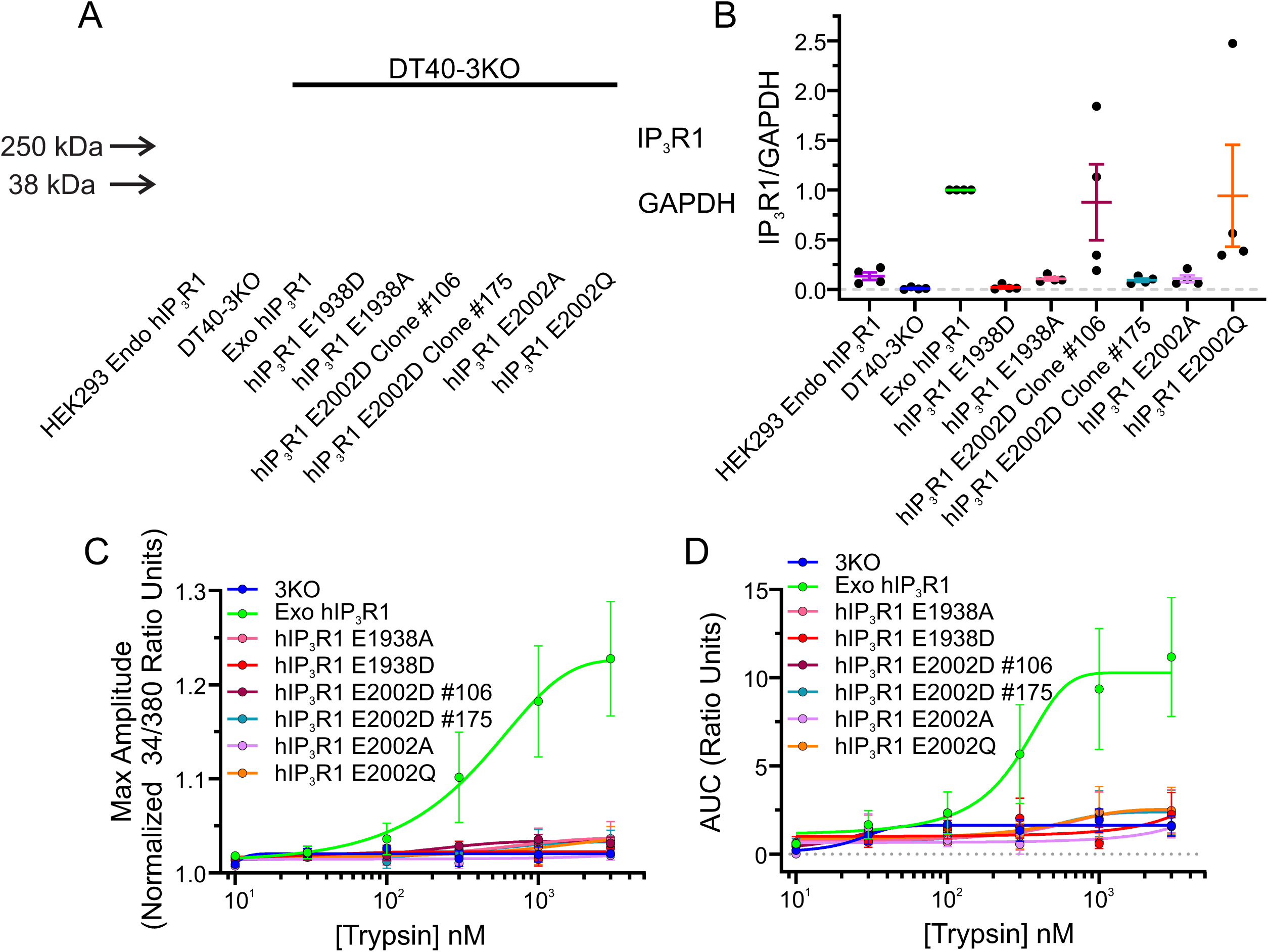
**A**. A representative western blot depicting IP_3_R1 and GAPDH protein levels in DT40-3KO, stable DT40-3KO over-expressing WT hIP_3_R1 (exo hIP_3_R1) and hIP_3_R1 with substitutions at the E1938 (E1938D and E1938A) and E2002 position (E2002D, E2002A and E2002Q). **B**. Scatter plot depicting quantification of IP_3_R1 protein level normalized to GAPDH from three-independent western blots. IP_3_R1/GAPDH expression level of all cell lines are normalized to that of exo hIP_3_R1. Dose-response curve showing **C**. maximum amplitude and **D**. AUC from Fura-2/AM loaded DT40-3KO (blue), exo hIP_3_R1 (green), hIP_3_R1 E1938A (pink), hIP_3_R1 E1938D (red), hIP_3_R1 E2002D (maroon, blue), hIP_3_R1 E2002A (purple) and hIP_3_R1 E2002Q (orange) cells in population-based assays when treated with increasing concentrations (100 nM, 300 nM, 1 μM, 3 μM, 10 μM, 30 μM and 100 μM) of trypsin. Data are mean ± SEM of three (N = 3) independent experiments.

**Figure S6.**
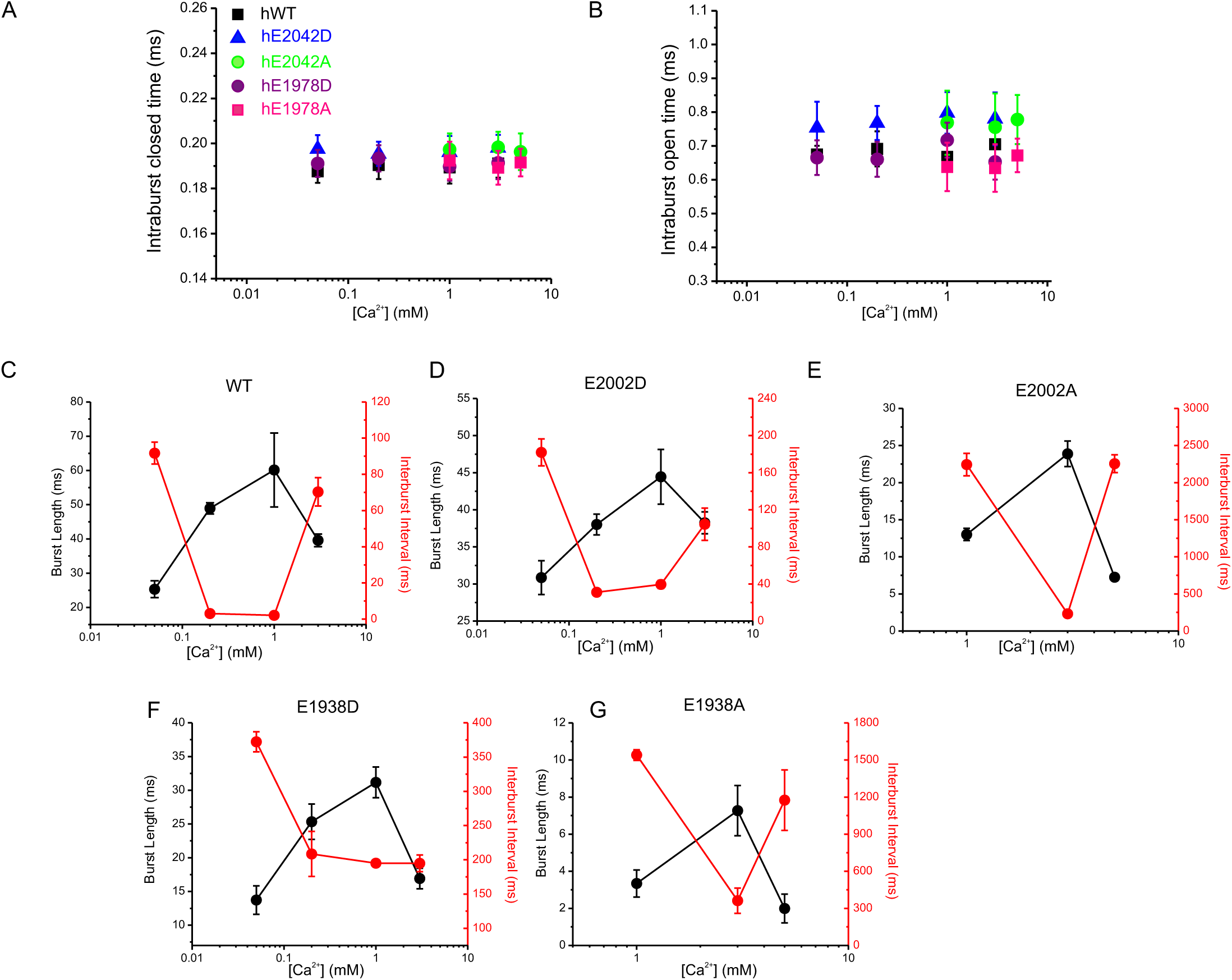
Analysis of intra and inter burst single channel open and closed times. **A**. Intraburst closed times for hIP_3_R1 and the indicated mutants are not altered. **B**. Intraburst open times for hIP_3_R1 and the indicated mutants are not altered. **C**. Burst length and interburst intervals for WT hIP_3_R with varying Ca^2+^. **D**. Burst length and interburst intervals for E2002A hIP_3_R1 with varying Ca^2+^. **E**. Burst length and interburst intervals for E2002D hIP_3_R1 with varying Ca^2+^. **F**. Burst length and interburst intervals for E1938D hIP_3_R1 with varying Ca^2+^. **G**. Burst length and interburst intervals for E21938A hIP_3_R1 with varying Ca^2+^. Increasing [Ca^2+^] decreases the interburst interval and increases the burst length at activating [Ca^2+^] and subsequently this relationship is reversed at higher [Ca^2+^] for all constructs.

---

**Table. S1.**
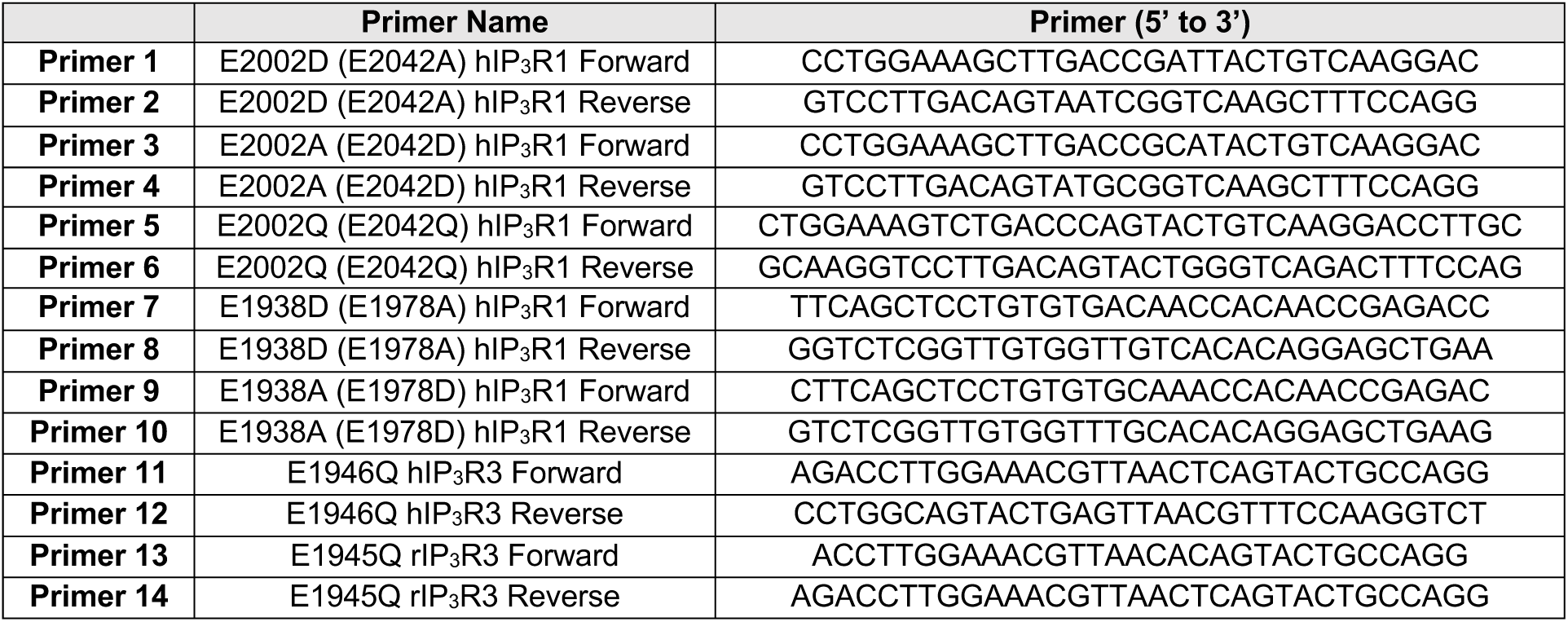
Primers used to generate substitutions at the desired site.

**Table. S2.**
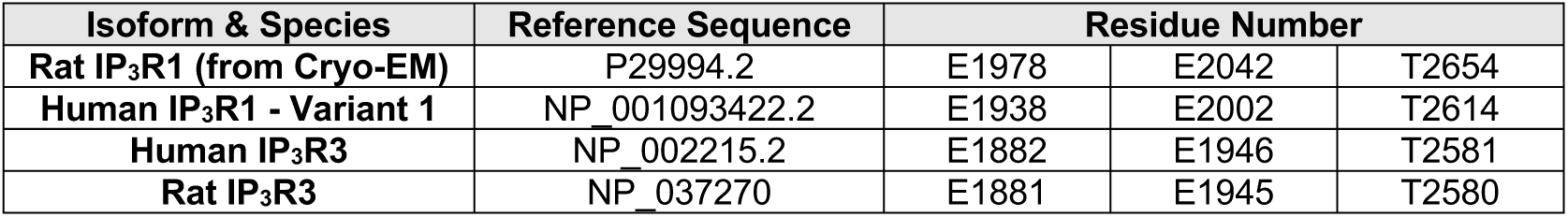
Reference sequence numbers and residues corresponding to Ca^2+^ binding residues.

## Notes

### Competing Interest Statement

The authors have declared no competing interest.

